# Nucleolar stress induced the formation of a nucleolar stress body via the NOSR-1/NUMR-1 axis in *Caenorhabditis elegans*

**DOI:** 10.1101/2024.03.05.583293

**Authors:** Minjie Hong, Xiaotian Zhou, Chenming Zeng, Demin Xu, Ting Xu, Shimiao Liao, Ke Wang, Chengming Zhu, Ge Shan, Xinya Huang, Xiangyang Chen, Xuezhu Feng, Shouhong Guang

## Abstract

Environmental stimuli not only alter gene expression profiles but also induce structural changes in cells. How distinct nuclear bodies respond to cellular stress is poorly understood. Here, we identified a new subnuclear organelle named the nucleolar stress body (NoSB), the formation of which was induced by the inhibition of rRNA transcription or inactivation of rRNA processing and maturation in *C. elegans*. NoSB did not colocalize with other previously described subnuclear organelles. We conducted forward genetic screening and identified a new bZIP transcription factor, named <u>n</u>ucle<u>o</u>lar <u>s</u>tress response-1 (NOSR-1), that is required for NoSB formation. The inhibition of rRNA transcription or inactivation of rRNA processing and maturation increased *nosr-1* expression. By using transcriptome analysis of wild-type animals subjected to different nucleolar stress conditions and *nosr-1* mutants, we identified that the SR-like protein NUMR-1 (nuclear localized metal responsive) is the target of NOSR-1. Interestingly, NUMR-1 is a component of NoSB and itself per se is required for the formation of NoSB. We concluded that the NOSR-1/NUMR-1 axis likely responds to nucleolar stress and mediates downstream stress-responsive transcription programs and subnuclear morphology alterations in *C. elegans*.

## Introduction

The nucleus contains several different dynamic bodies that are variously composed of proteins or specific RNA molecules (1). Intriguingly, nuclear bodies are not surrounded by membranes yet are retained as separate and distinct entities within the nucleoplasm, formed by a process termed liquid–liquid phase separation (LLPS) (2, 3). It is well established that LLPS-associated proteins that contain folded domains, intrinsically disordered regions (IDRs) and low complexity (LC) regions play vital roles in the process of nucleation (4). For instance, Paraspeckles require NONO, PCPC1, and PSF (5, 6). Cajal bodies are disrupted or disappear in the absence of COILIN, SMN, FAM118B, or WRAP53 (7, 8), PML bodies are nucleated by PML (9), and SON and SRRM2 are essential for nuclear speckle formation (10).

Nuclear body formation is generally dynamic, especially in response to a variety of stress conditions (11–13). For example, heat shock, UV radiation, and chemical toxicity trigger the formation of nuclear stress bodies (nSBs), which may further influence the reprogramming of gene expression (14). Mitochondrial stress leads to altered morphology and numbers of nuclear paraspeckles (15). Polycomb Group (PcG) bodies were shown to repress damaged DNA upon UV irradiation (16). However, how these nuclear bodies respond to cellular stresses and their physiological roles are still poorly understood.

The nucleolus is the most prominent subnuclear structure, where the main processes of rRNA and ribosome biogenesis occur. In the mammalian nucleolus, rDNA genes are transcribed by RNA polymerase I into 47S pre-rRNA, which is subsequently processed and modified to generate 28S, 18S, and 5.8S rRNAs. Genes encoding 5S rRNA are transcribed by Pol III (17, 18). These rRNAs are assembled with ribosomal proteins (RPs) to form small and large preribosome subunits, which are exported to the cytoplasm and eventually form the mature 40S and 60S ribosome subunits (19).

Ribosome assembly factors maintain their greatest steady-state concentrations within nucleoli under normal growth conditions (20), and blocking ribosome biosynthesis stimulates nucleolar stress (21). For instance, actinomycin D inhibits Pol I transcription elongation and causes nucleolar segregation (22). Starvation or rapamycin treatment downregulates mTOR signaling in yeast and mammalian cells, leading to a significant nucleolar size reduction and inhibition of rRNA transcription (23). Mutations in ribosome assembly factors or ribosomal proteins (r-proteins) lead to nucleolar stress and induce a number of human disorders (24).

In mammals, nucleolar stress signaling pathways are classified into two types: p53-dependent and p53-independent pathways (20). p53 is a crucial transcription factor that responds to many cellular stresses. Nucleolar stress releases p53 from inhibition by MDM2 and activates a set of target genes (25, 26). The mechanism of the p53-independent pathway remains poorly understood. Nucleolar stress may induce cell proliferation or cell cycle arrest via the regulation of c-Myc, E2F-1, p21 Waf1/Cip1 or p27Kip1, independent of p53 (27–30). Yeast and plants do not express p53 or MDM2 and therefore use mechanisms independent of the p53 pathway. In plant cells, the plant-specific transcription factor ANAC082 may play critical roles in the response to nucleolar stress (31).

*C. elegans* expresses p53/CEP-1 but lacks MDM2. Active p53/CEP-1 is regulated at the translational level (32, 33). Mutation in the conserved nucleolar protein NOL-6 enhances resistance to bacterial infection by activating p53/CEP-1 and its target gene, SYM-1 (34). The depletion of a nucleolar protein, WDR-46, activates xenobiotic detoxification genes via Nrf/SKN-1 and p53/CEP-1 (35). Interestingly, the transcription factor PHA-4/FoxA, instead of CEP-1/P53, acts as a nucleolar stress sensor to induce the expression of lipogenic genes and lipid accumulation (36), suggesting additional mechanisms for the nucleolar stress response.

Previously, we reported that nucleolar stress could induce the accumulation of erroneous rRNAs that lead to the recruitment of RdRPs to synthesize antisense ribosomal siRNAs (risiRNAs), which subsequently turn on the nuclear RNAi-mediated gene silencing pathway to inhibit pre-rRNA expression (28, 54, 55). In addition, EXOS-10 exhibits nucleoplasmic translocation upon cold-worm shock in *C. elegans* (56). Third, abnormal accumulation of 27SA_2_ rRNA intermediates upon the depletion of class I RPLs reshaped spherical nucleoli to a ring-shaped nucleolar structure with an enlarged nucleolar vacuole (NoV) in *C. elegans* (26). Therefore, it is very likely that a number of cellular events may occur concurrently to respond to nucleolar stress.

Here, we identified that nucleolar stress induced the formation of a new subnuclear organelle, named the nucleolar <u>s</u>tress <u>b</u>ody (NoSB), in *C. elegans*. The inhibition of rRNA transcription by actinomycin D, inactivation of rRNA processing and maturation, or mutation in the exosome ribonuclease complex, which degrades erroneous rRNAs, results in NoSB formation. To search for factors required for the formation of nucleolar stress-induced NoSB, we conducted forward genetic screening and identified a new bZIP-containing protein, <u>n</u>ucle<u>o</u>lar <u>s</u>tress <u>r</u>equired-1 (NOSR-1). In the absence of NOSR-1, nucleolar stress failed to induce NoSB formation. Interestingly, NOSR-1 acts as a sensor of nucleolar stress to activate the expression of the SR-like protein NUMR-1, consequently promoting NoSB formation. Therefore, our work revealed a novel NOSR-1/NUMR-1-involved pathway that responds to nucleolar stress and mediates nuclear substructure alterations in *C. elegans*.

## Results

### Deficiency in rRNA processing and maturation induced the formation of a novel nucleolar stress body

Our recent work showed that the depletion of class I RPL proteins and a number of 26S rRNA processing factors led to nucleolar reshaping, in which the 27SA_2_ rRNA intermediate accumulated, nucleolar rings formed, and nucleolar vacuoles enlarged (Fig. 1A) (37). To further investigate the nucleolar stress-induced substructure alterations in the nucleus, we generated an mCherry::DIS-3 strain and visualized the structure of the nucleus and nucleolus in hypodermal cells upon a number of nucleolar stresses. DIS-3 (also known as Rrp44 or EXOSC11) is a subunit of the 3’ to 5’ exoribonuclease complex, which degrades erroneous rRNAs in the nucleus and accumulates in the nucleoplasm (Fig. 1B) (38, 39). The single copy mCherry::DIS-3 transgene was integrated onto the *C. elegans*’ chromosome III of the strain EG8080 by MosSCI technology (39). The transgene was driven by its own promoter, did not affect the brood size, but rescued *dis-3(ust56)* dysfunction in risiRNA biogenesis, suggesting that the transgene recapitulated the function of endogenous DIS-3 (39).

**Figure 1.**
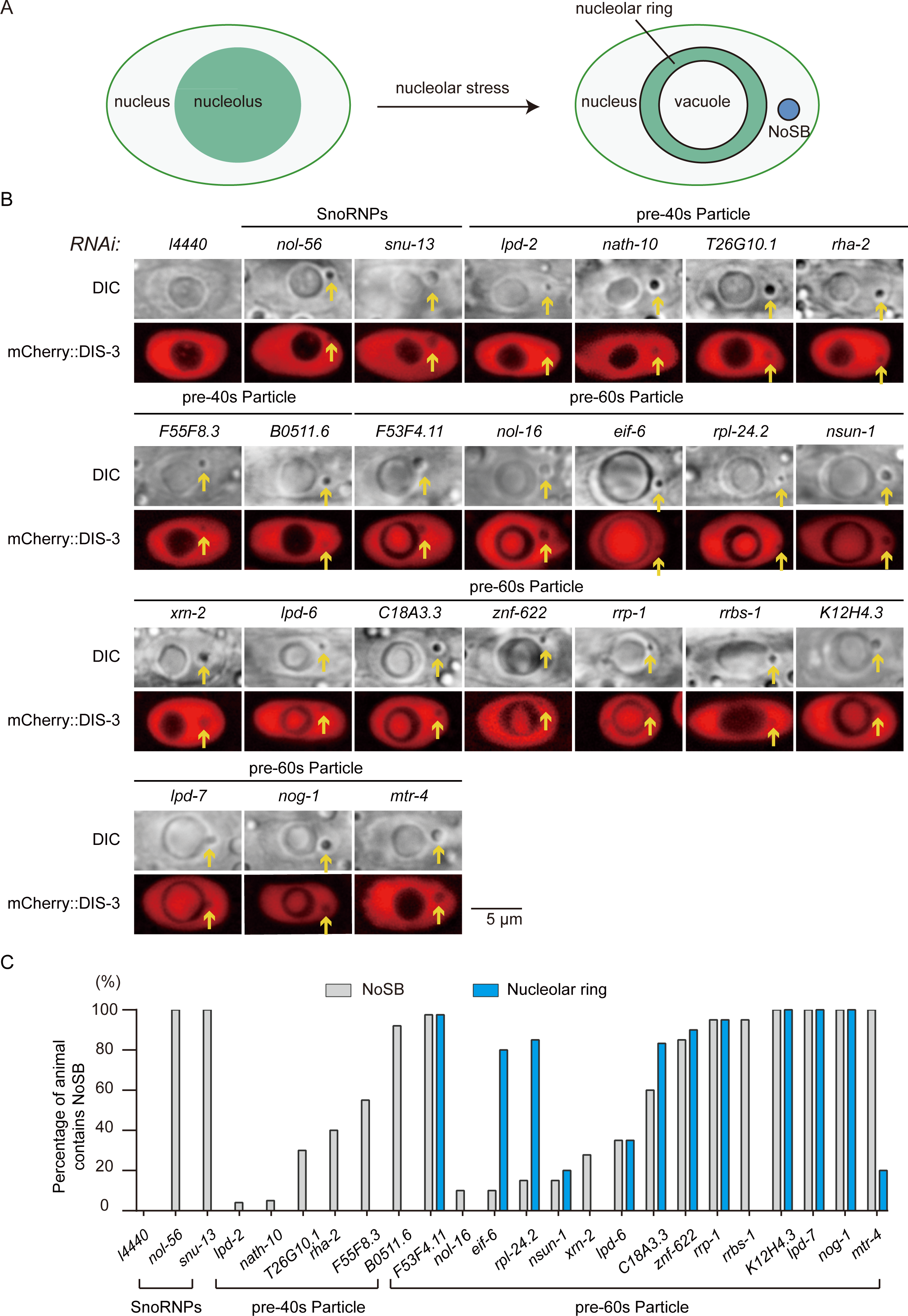
Deficiency in rRNA processing and maturation induced the formation of a novel nucleolar stress body. (A) Schematic diagram of the formation of nucleolar rings, enlarged nucleolar vacuoles and nucleolar stress bodies (NoSBs) in *C. elegans*. (B) DIC and fluorescence microscopy images of nuclei in hypodermal cells after knockdown of the indicated genes in the indicated animals. (C) Quantification of NoSB and ring-shaped nucleoli after RNAi knockdown of the indicated genes. The percentage of animals containing at least three NoSB structure were shown. n>19 animals.

We performed feeding RNAi experiments and knocked down approximately 100 genes involved in rRNA processing and modification in mCherry::DIS-3 animals (Table S1). Then, we visualized the hypodermal cells using Nomarski (DIC) and fluorescence microscopy. Consistent with previous work, we observed the formation of nucleolar ring structures and enlarged nucleolar vacuoles upon knocking down a number of genes involved in pre-60S ribosome maturation (Figs. 1B-C) (37). The nucleolar vacuole (NoV) is an evolutionarily conserved nucleolar subcompartment, presents in the nucleoli of various plants and animals (40–43). Our recent work showed that faithful rRNA processing is essential to maintain the structure of nucleolus (37). When the rRNA precursor 27SA_2_ rRNA was inappropriately accumulated, the spherical nucleolus transformed into ring-shaped nucleolus, in which a NoV appears. In the circumstance, the rRNA transcription and processing machineries were enriched in the ring, while many usual nucleoplasmic proteins accumulated in the NoV (37). NoVs may have important roles in germline development or mRNA metabolism and may be involved in the transport of nucleolar substance from nucleolus to nucleoplasm and temporary storage of certain materials.

Strikingly, we noticed the appearance of an additional vacuole per cell in the nucleoplasm in hypodermal cells (Figs. 1B-C). The vacuole excludes DIS-3 and is also visible by DIC microscopy. We named this vacuole the <u>n</u>ucle<u>o</u>lar <u>s</u>tress <u>b</u>ody (NoSB) (see below).

Ribosomal proteins of the small subunit (RPS) fall into two main categories according to their functions in pre-rRNA processing (44). i-RPSs are strictly required for initiating the processing of sequences flanking the 18S rRNA. p-RPS proteins are required for subsequent nuclear and cytoplasmic maturation steps. Knocking down both i-RPS and p-RPS proteins induced the formation of NoSB but not the formation of nucleolar rings (Figs. 1B, 1C, S1A, S1C) (37).

RPLs are proteins of the 60S ribosome large subunit that are involved in pre-rRNA processing, including 27SA_2_, 27SA_3_, 27SB pre-rRNAs and 7S pre-rRNA maturation. (45). *C. elegans*’ genome encodes 48 large subunit (*rpl*) genes. Our recent work classified RPL proteins into two main functional categories according to whether they are required for the processing of 27SA_2_ pre-rRNAs and the suppression of ring-shaped nucleoli and enlarged nucleolar vacuoles (NoV) (37). The Class I RPLs are required for 27SA_2_ pre-rRNA processing and knockdown this group significantly increased the proportion of vacuole-contained nucleoli and the accumulation of 27SA_2_ pre-rRNA (37). However, Class II RPLs were likely involved in the processing of other pre-rRNA intermediates, and knockdown of Class II RPLs did not lead to the formation of NoV (37). The depletion of both classes of RPL proteins induced NoSB formation, yet only class I RPLs triggered the formation of ring-shaped nucleoli (Figs. S1B-C).

Taken together, these results suggested that defective rRNA biogenesis and ribosome assembly could induce alterations in subnuclear structures in *C. elegans*.

### Mutation of exosome ribonucleases induced NoSB formation

To investigate the mechanism of NoSB formation, we first conducted forward genetic screening to search for mutants that reveal NoSB formation. We mutagenized the mCherry::DIS-3 strain with ethyl methanesulfonate (EMS) followed by clonal screening with DIC and fluorescence microscopy (Fig. 2A). From the 2000 haploid genome, we isolated two mutants, *ust242* and *ust243*. Using whole-genome resequencing, we identified two point mutations in the coding regions of *exos-4.2 (ust243, G54R)* and *exos-10 (ust242, D368N)* (Fig. 2B). Both EXOS-4.2 and EXOS-10 are core factors of the exosome 3’-5’-exoribonuclease complex (38, 39, 46). *exos-4.2* encodes a ribonuclease (RNase) pleckstrin homology (PH)-like protein, and *exos-10* encodes a 3’-5’-exoribonuclease.

**Figure 2.**
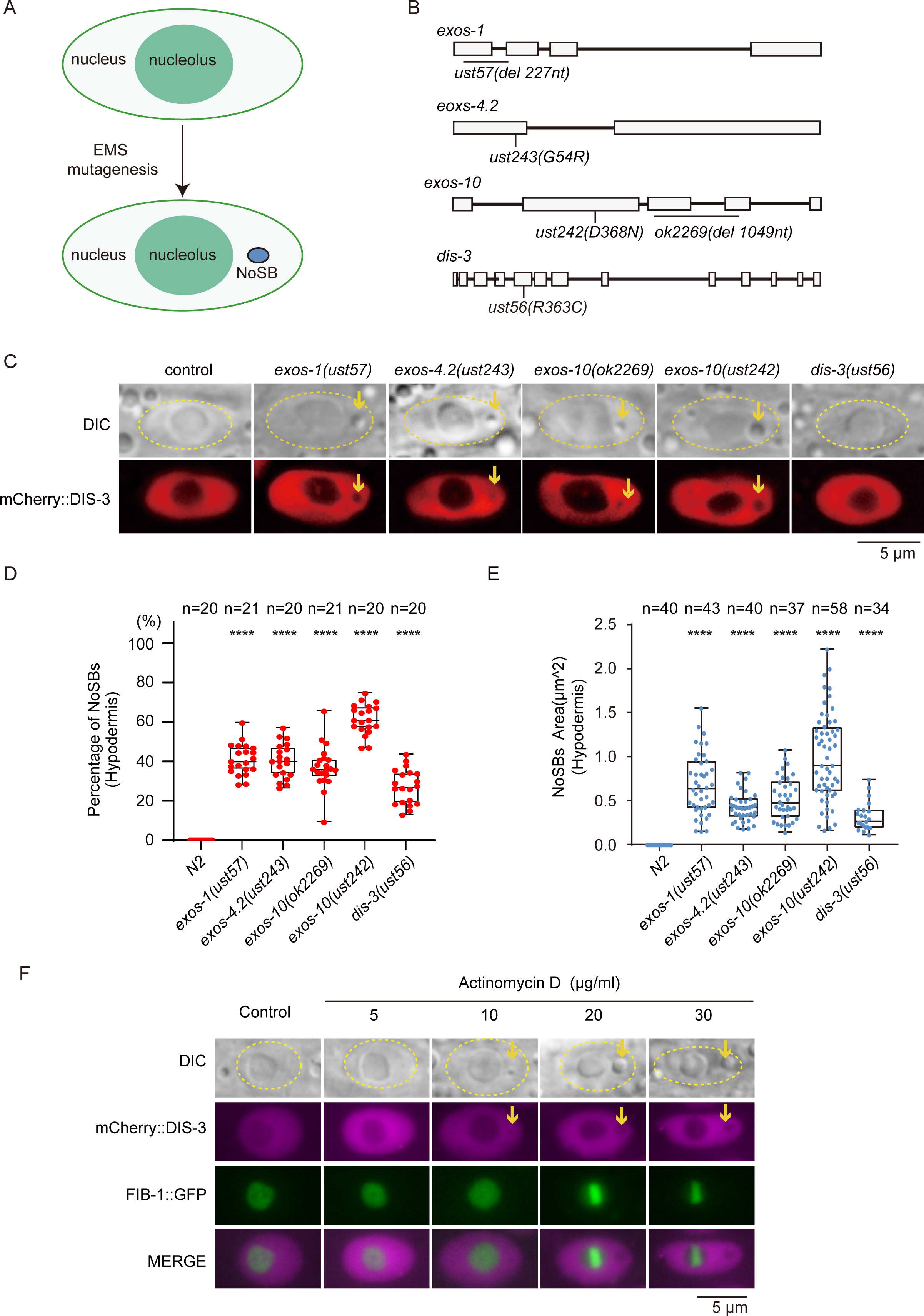
Mutation of exosome ribonucleases induced NoSB formation. (A) Schematic diagram of forward genetic screening to search for factors suppressing NoSB formation in *C. elegans* nuclei. (B) Schematic of the gene structure of exosome genes. *exos-4.2 (ust243*) and *exos-10 (ust242)* are the mutants obtained from the forward genetic screen. *exos-1 (ust57), exos-10 (ok2269)* and *dis-3 (ust56)* were described previously (39, 90). (C) DIC and fluorescence microscopy images of nuclei in hypodermal cells in the indicated animals. (D) Quantification of NoSB in hypodermic cells in exosome mutants. Every dot indicates the percentage of hypodermis cells containing NoSB in each worm. n>19 animals. (E) The size of NoSB in exosome mutants. n>30 nucleoli from n>19 animals. (F) DIC and fluorescence microscopy images of *C. elegans* nuclei in hypodermal cells after actinomycin D treatment.

To confirm that the exosome complex suppresses NoSB formation, we generated additional alleles of *exos-4.2* and *exos-10* by CRISPR/Cas9 technology and obtained mutants of other exosome subunits. The mutation of *exos-1, exos-4.2* and *exos-10* resulted in NoSB formation (Figs. 2C-E). Consistently, feeding mCherry::DIS-3 animals dsRNAs targeting *exos-2*, *exos-3*, *exos-4.1*, *exos-5*, *exos-7, exos-8*, or *exos-9* induced the formation of NoSB (Fig. S2A).

We measured the size of NoSB and nucleoli in the exosome mutants. The size of NoSB is approximately 10% of that of nucleoli, and the exosome mutants exhibited modestly enlarged nucleoli (Figs. 2E and S2B). Similarly, in *S. cerevisiae* and *D. melanogaster,* the depletion of ribosomal proteins also led to enlarged nucleoli (47).

LMN-1 is an ortholog of human lamin A/C and localizes to the inner side of the nuclear membrane. NoSB localized inside the nuclei, as shown by LMN-1::mCherry (Fig. S2C). MTR-4 and NRDE-3 are two nucleoplasm-localized proteins and are involved in nuclear and nucleolar RNAi (39). Our previous work showed that deficient rRNA processing and maturation led to the accumulation of erroneous rRNAs and induced the production of a class of antisense ribosomal siRNA (risiRNA) which further translocated the Argonaute protein NRDE-3 from the cytoplasm to nucleoli (39, 48). Similarly, the disruption of RNA exosomes also resulted in the production of risiRNAs and triggered the nucleolar accumulation of NRDE-3 (39). To test whether nucleolar stress could also induce the production of risiRNAs and enrich NRDE-3 in NoSBs, we visualized the subcellular localization of NRDE-3 and MTR-4 upon *exos-9* RNAi. However, both proteins were excluded from the *exos-9* depletion-induced NoSB (Fig. S2D), suggesting that it is unlikely risiRNA could enrich in NoSB.

Therefore, we concluded that a deficient exosome ribonuclease complex could induce NoSB formation.

### The inhibition of RNA polymerase I induced NoSB formation

Previous work showed that actinomycin D inhibits rRNA transcription and induces nucleolar stress (37, 49, 50). We generated an mCherry::DIS-3;FIB-1::GFP strain and visualized the structure of the nucleolus in hypodermal cells in the presence of actinomycin D. FIB-1 encodes the *C. elegans* ortholog of human fibrillarin and *Saccharomyces cerevisiae* Nop1p (51) and localizes to the nucleolus.

At a concentration of 5 µg/ml actinomycin D, we did not observe pronounced nuclear and nucleolar substructure changes in hypodermal cells (Figs. 2F and S3A). However, at a higher concentration of 20 to 30 µg/ml actinomycin D, FIB-1 likely condensed and detached from some unidentified nucleolar border. The NoSB excludes DIS-3 in the nucleoplasm, which is also visible by DIC and fluorescence microscopy. FIB-1 did not accumulate in NoSB, suggesting that the vacuole is distinct from nucleoli. The sizes of NoSB increased at a higher concentration of actinomycin D (Fig. S3B). LMN-1::mCherry confirmed that actinomycin D-induced NoSB localized in the nucleus (Fig. S3C). MTR-4 and NRDE-3 were also excluded from actinomycin D-induced NoSB (Fig. S3D). Notably, actinomycin D treatment has been shown to inhibit the formation of nucleolar ring and nucleolar vacuole (37), which is oppositive to actinomycin D-induced NoSB formation.

We tested whether other kinds of environmental conditions could induce the formation of NoSB. However, growing animals at 15°C or 25°C, cold shock at 4°C or heat shock at 37°C failed to induce the formation of NoSB (Figs. S3E-G, also see Figs. S18A-B). In addition, the ectopic expression of mcherry::DIS-3 did not affect the efficacy of NoSB formation (Figs. S4A-B).

Taken together, these data suggested that the inhibition of RNA polymerase I by actinomycin D induced the formation of NoSBs.

### NoSB is not a portion of nucleoli that blebbed off upon nucleolar stress

In mammalian cells, the factors involved in rRNA transcription and processing and ribosome assembly usually localize to the FC, DFC, and GC nucleolar subcompartments. Recent work showed that *C. elegans* nucleoli contain two phase-separated subcompartments, with FIB-1 and GARR-1 in the FC and NUCL-1 and LPD-6 in the GC region (40).

To test the possibility that NoSB may be a portion of the nucleolus that has blebbed off upon nucleolar stress, we generated a number of single-copy transgenes of nucleoli-localized proteins by CRISPR-Cas9 technology (Figs. 3A-C, S5A). Among them, FIB-1 is a methyltransferase for pre-rRNA processing and modification. RBD-1 is an ortholog of human RBM19, which is required for 90S pre-ribosome maturation. RRP-8 is a rRNA processing factor involved in N1-methyladenosine (m1A) modification of 26S rRNAs. DAO-5 is an rRNA transcription factor. RPL-7 and T06E6.1 are rRNA processing factors. GARR-1 involved in snoRNA guided rRNA pseudouridine synthesis. RPOA-1, RPOA-2, and RPAC-19 are subunits of the RNA polymerase I complex. We observed that upon *exos-9* RNAi, all these proteins were excluded from the NoSB (Figs. 3B, 3C, S5A, and summarized in Fig. S5B).

**Figure 3.**
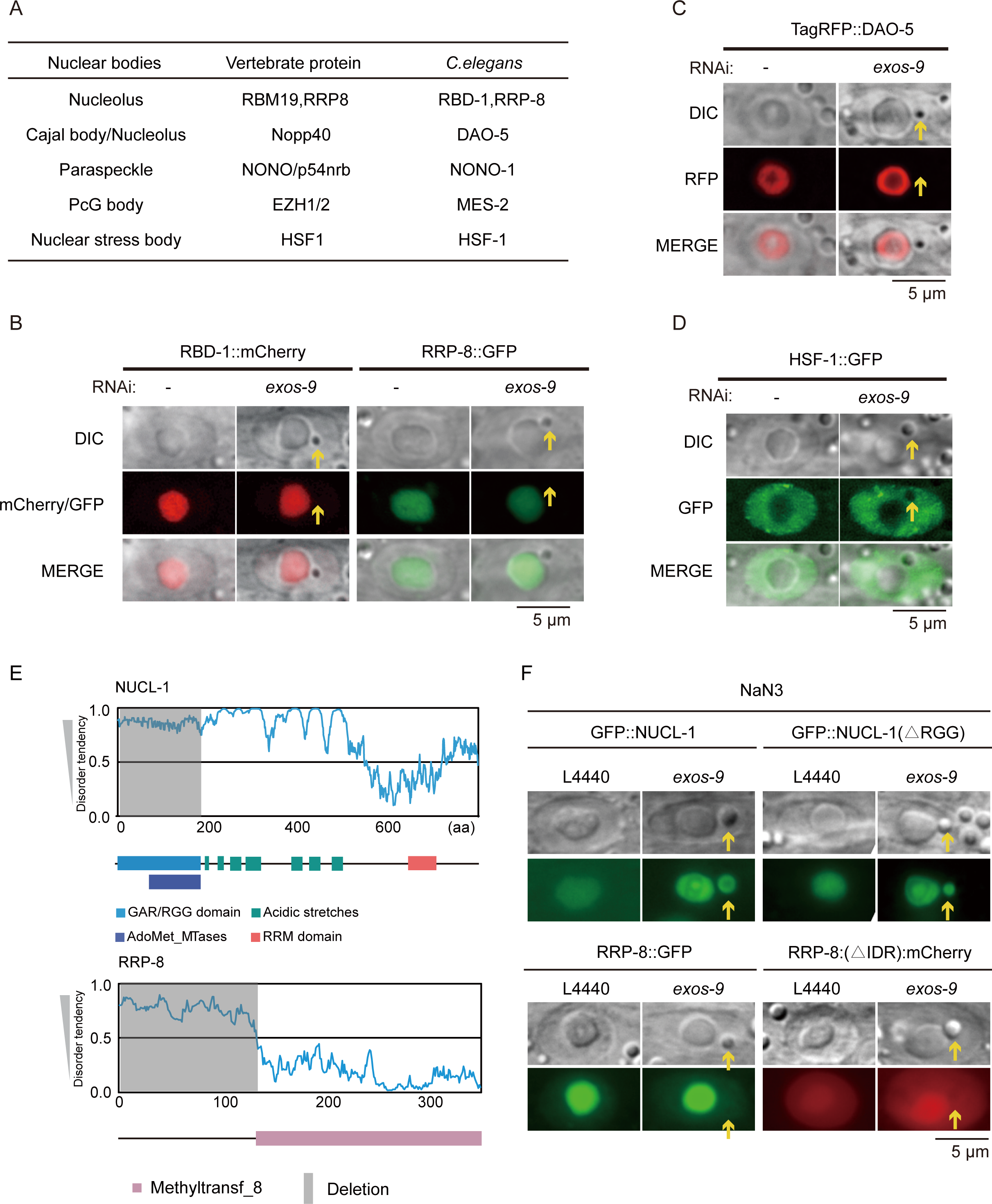
NoSB is a new subnuclear organelle. (A) List of marker genes of subnuclear organelles and their *C. elegans* homologs. (B-D) DIC and fluorescence microscopy images of nuclei with the indicated transgenes after *exos-9* knockdown. (E) Schematic diagram of the domain structure and predicted intrinsically disordered regions of NUCL-1 and RRP-8. (F) DIC and fluorescence microscopy images of nuclei with the indicated transgenes after *exos-9* knockdown.

Therefore, we concluded that NoSB is not a portion of the nucleolus that has blebbed off upon nucleolar stress.

### NoSB is a new subnuclear organelle

To test whether NoSB is a new subnuclear organelle or colocalizes with other known subnuclear structures, we generated a number of fluorescence-labeled protein markers to reveal the Cajal body, paraspeckle, PcG body and nuclear stress body (Fig. 3A) (52). The *exos-9* depletion-induced NoSB did not colocalize with any of these known subnuclear organelles (Figs. 3D, S6A-D). The heat shock protein HSF-1 responds to heat stress by forming subnuclear structures termed nuclear stress granules in *C. elegans*. However, we did not find colocalization of NoSB with HSF-1::GFP (Fig. 3D).

We tested whether NoSB is a nuclear lipid droplet. Recent work reported that lipid droplets, which form at the nuclear envelope, may enter the nucleoplasm by penetrating the nuclear lamina and associated peripheral heterochromatin to form nuclear lipid drops (nLDs) (53, 54). Actinomycin D-triggered nucleolar stress could induce excessive lipid accumulation in *C. elegans* (36). To test whether NoSB is a kind of nLD, we used Oil Red O to stain neutral lipids of postfixed animals with or without *exos-9* RNAi but failed to observe lipid accumulation in NoSBs (Fig. S6E). Therefore, NoSB is unlikely to form nuclear neutral lipid droplets.

An interestingly observation is NUCL-1. NUCL-1 located in GC region and is required for nucleolar vacuole formation (37). NUCL-1 encodes an evolutionarily conserved protein exhibiting extensive homology to yeast and human nucleolin and has been shown to interact with exosome proteins (55). The N-terminal domain of NUCL-1 is a long IDR containing a GAR/RGG domain that is required for subnucleolar compartmentalization (40) (Fig. 3E). IDRs are likely one of the driving forces of phase separation and have been identified in many proteins capable of phase separation. RRP-8 encodes an evolutionarily conserved protein exhibiting extensive homology to yeast and human RRP8. The N-terminal domain of RRP-8 is also a long IDR (Fig. 3E). We constructed GFP- or mCherry-tagged NUCL-1, NUCL-1ΔRGG, RRP-8, and RRP-8Δ IDR transgenic animals. The depletion of RNA exosome did not translocate NUCL-1 or NUCL-1(ΔRGG) into NoSB in live worms (Figs. S7B-C). However, a further treatment with NaN_3_ following Actinomycin D or *exos-9* RNAi could induce an accumulation of NUCL-1 or NUCL-1(ΔRGG) to NoSB (Figs. 3F, S7A-C). Yet *nucl-1* is not required for the formation of NoSB (Fig. S7D). The reason is not clear yet. We suspected that NUCL-1 per se is not a physiological component of NoSB, and NaN_3_-induced hypoxia or cell death might further change the biochemical properties or binding partners of NUCL-1 that redistributed the protein to NoSB upon nucleolar stress. Neither RRP-8 nor RRP-8(ΔIDR) translocated to NoSB upon nucleolar stress followed by NaN_3_ treatment (Fig. 3F).

In conclusion, we speculated that NoSB is a new subnuclear structure induced by nucleolar stress.

### Forward genetic screening identified a bZIP transcription factor NOSR-1 required for NoSB formation

To further understand the mechanism and regulation of NoSB, we performed a second forward genetic screening to search for factors that were required for NoSB formation. We chemically mutagenized mCherry::DIS-3;*exos-10(ust242)* animals followed by clonal screening with DIC and fluorescence microscopy and isolated mutants in which NoSB was depleted (Fig. 4A). From approximately 2000 haploid genomes, we isolated one mutant, *ust303*, in which NoSB was depleted in mCherry::DIS-3;*exos-10(ust242)* animals (Fig. 4B). We named the gene <u>n</u>ucle<u>o</u>lar <u>s</u>tress response-1 (*nosr-1*).

**Figure 4.**
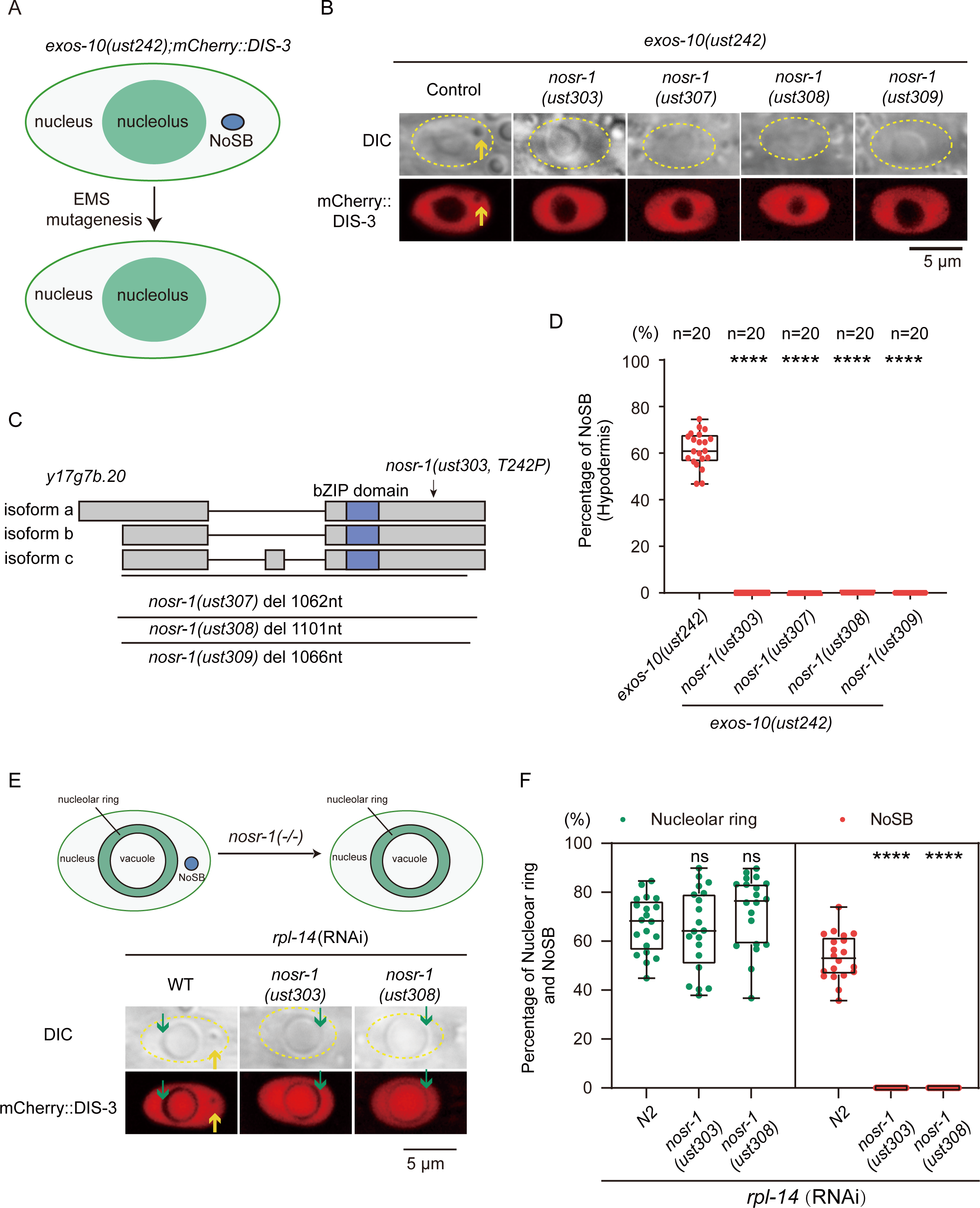
Forward genetic screening identified a bZIP transcription factor NOSR-1 required for NoSB formation. (A) Schematic diagram of the forward genetic screening to search for factors required for NoSB formation in *C. elegans* nuclei. (B) DIC and fluorescence microscopy images of nuclei in the indicated animals. (C) Schematic of the *nosr-1* exon structure. The deletion alleles *ust307*, *ust308* and *ust309* were constructed by CRISPR/Cas9 technology. (D) Quantification of NoSB in hypodermic cells in the indicated animals. Every dot indicates the percentage of hypodermis cells containing NoSB in each worm. n>19 animals. (E) Images of mCherry::DIS-3 transgene in *nosr-1*(*-*)*;rpl-14(RNAi)* animals. (F) Quantification of nucleolar rings and NoSB in hypodermic cells in the indicated animals. Every dot indicates the percentage of hypodermis cells containing NoSB in each worm.

We deep-sequenced the *nosr-1(ust303)* genome and identified Y17G7B.20 (Fig. 4C), in which the amino acid threonine242 was mutated to a proline (Fig. 4C). Y17G7B.20 was predicted to have a basic leucine zipper (bZIP) domain that can bind to DNA and mediate protein dimerization (56, 57). To further confirm that *y17g7b.20* is *nosr-1*, we generated three additional deletion alleles of *y17g7b.20* by CRISPR/Cas9 technology with two single-guide RNAs (sgRNAs) (58) and examined NoSB formation in *exos-10(ust242)* animals. All three alleles, *ust307, ust308* and *ust309*, deleted most of the gene sequence and caused a frame shift; therefore, they are likely null alleles. In the three *y17g7b.20* mutants, *exos-10(ust242)* mutation or RNAi knockdown of *exos-9* failed to induce NoSB formation (Figs. 4B, 4D, and S8A-B). In addition, the *nosr-1* mutation blocked actinomycin D-induced NoSB formation (Fig. S8C). Therefore, *y17g7b.20* is *nosr-1*.

To test whether NOSR-1 is the key player in NoSB formation from all types of defects in ribosome synthesis, we observed that (1) *nosr-1* is required for both *exos-9* RNAi- and Actinomycin D-induced NoSB formation (Figs. S8A-C); (2) knockdown the components of snoRNP, pre-40S and pre-60S particles, and RPSs and RPLs induced NoSB formation, which also depended on *nosr-1* (Figs. S9A-B, S10A-B). These data suggested that NOSR-1 likely suppresses the formation of NoSB induced by all types of defects in ribosome biogenesis.

Interestingly, the depletion of *nosr-1* inhibited *rpl-14* knockdown-induced NoSB formation but did not prohibit the formation of ring-shaped nucleoli (Figs. 4E-F), suggesting that the formation of NoSB and ring-shaped nucleoli involved two distinct mechanisms.

### NOSR-1 promotes animal fertility and lifespan under nucleolar stress conditions

The bZIP transcription factor family is a gene family conserved from yeast to humans and has roles in a wide variety of processes (59). *C. elegans* underwent a relatively recent expansion of bZIP genes, some of which do not appear to have direct homologs in mammals, likely including NOSR-1 (56, 59).

We investigated the role of NOSR-1 in *C. elegans*. *nosr-1* mutants revealed brood sizes similar to those of wild-type N2 animals at 20°C but strongly reduced fertility at 25°C (Figs. 5A-B). In addition, *nosr-1* mutation reduced the brood size of *exos-10* animals (Figs. 5C-D).

**Figure 5.**
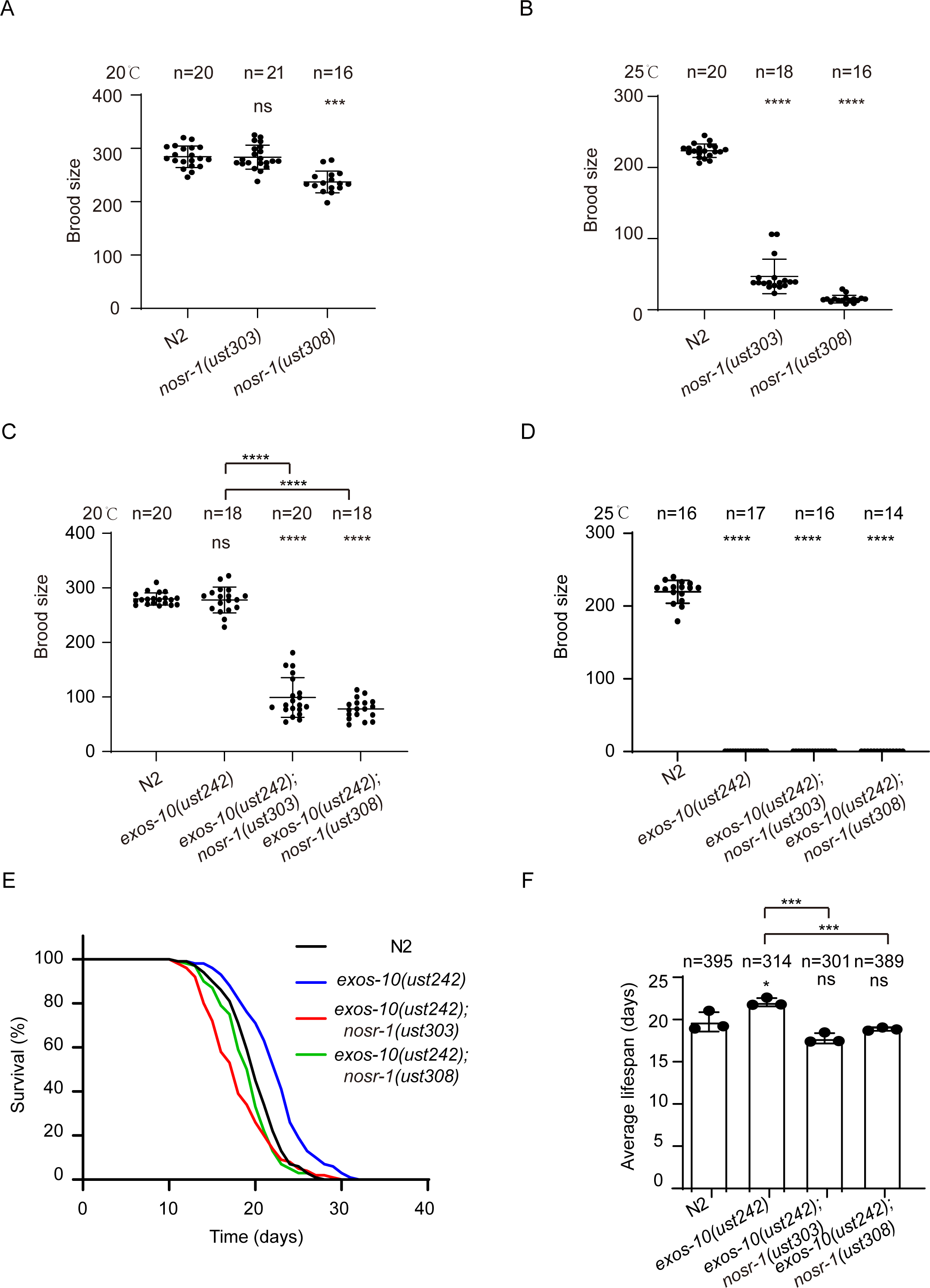
NOSR-1 is required for fertility and lifespan. (A-D) Brood size of the indicated animals at 20°C and 25°C. (E) Survival curves of the indicated animals at 20°C. (F) Histogram displaying the average lifespan of the indicated animals. Mean ± SD of three independent experiments. Asterisks indicate significant differences using log rank tests. ns, not significant, p>0.05.

We failed to detect a significant change in the lifespan of *nosr-1* animals compared to that of N2 animals (Figs. S11A-B). However, the *nosr-1* mutation shortened the lifespan of *exos-10* animals (Figs. 5E-F). The depletion of *nosr-1* did not significantly change the oxidative stress resistance or heat stress resistance in N2 background worms (Figs. S11C–D).

Therefore, we concluded that *nosr-1* is required for fertility and lifespan regulation in animals under nucleolar stress conditions.

### Nucleolar stress activates *nosr-1* expression

To investigate the mechanism of *nosr-1* in NoSB regulation, we tested whether *nosr-1* responds to nucleolar stresses. We compared the transcriptomes of animals with or without nucleolar stress using mRNA deep sequencing. The *nosr-1* mRNA levels were upregulated upon actinomycin D treatment, *exos-10* mutation, or RNAi knockdown of *exos-9* and *rpl-14* (Fig. 6A).

**Figure 6.**
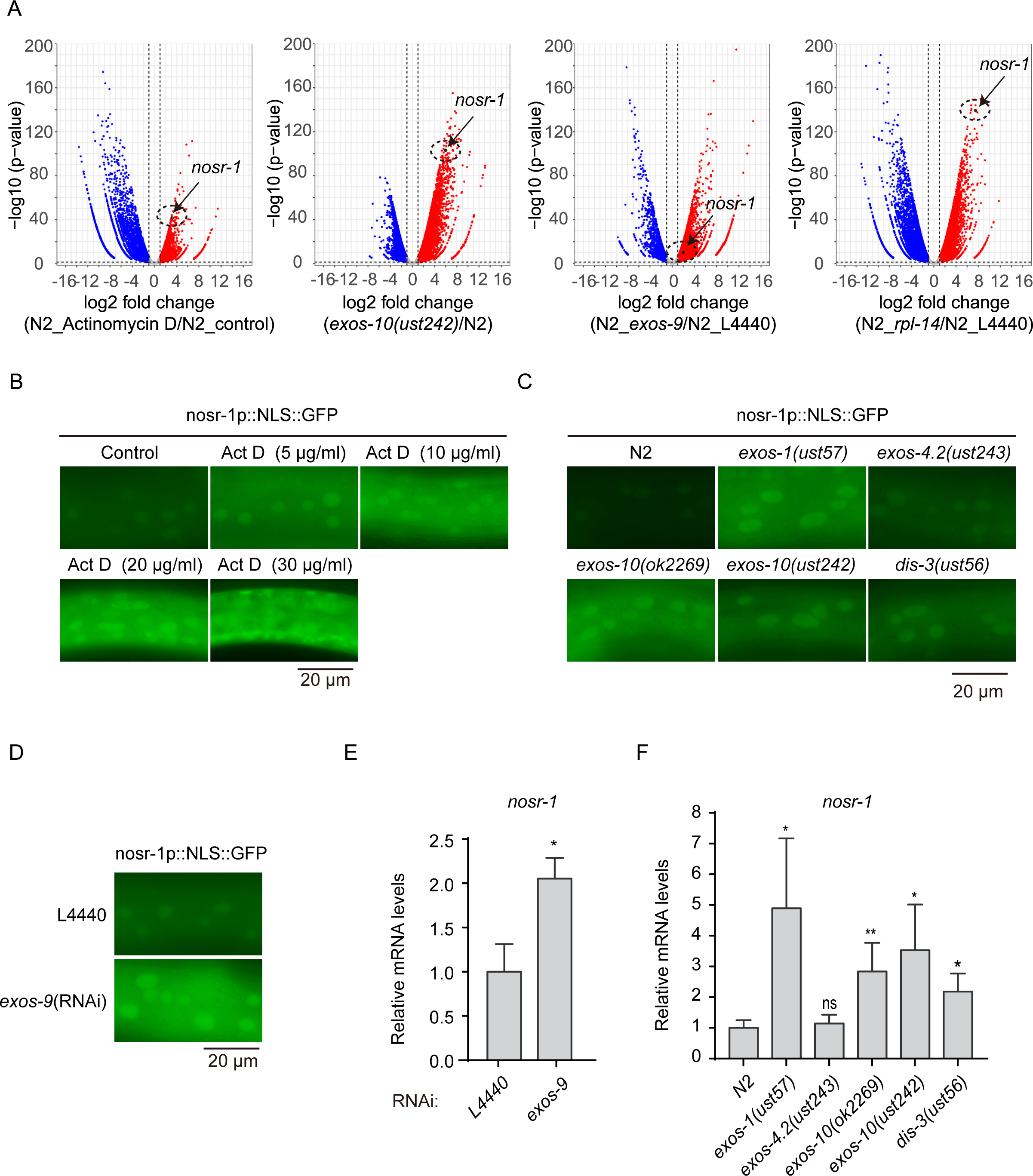
Nucleolar stresses activate *nosr-1* expression. (A) Volcano plot comparing the transcriptomes of the indicated animals. The expression of *nosr-1* is highlighted. (B) Fluorescence microscopy images of *nosr-1_p_::NLS::GFP(ustIS285)* after actinomycin D treatment. (C) Fluorescence microscope images of *nosr-1_p_::NLS::GFP(ustIS285)* in the indicated animals. (D) Fluorescence microscope images of *nosr-1p::NLS::GFP* after knocking down the indicated genes by RNAi. (E-F) Relative mRNA levels of *nosr-1* measured by quantitative real-time PCR in the indicated animals. Data are presented as the means ± SD of three biological replicates. ****P < 0.0001, **P < 0.01, *P < 0.05.

Then, we constructed a GFP-fused transcription fusion *nosr-1_P_::NLS::GFP* and inserted it at the ttTi4348 site on chromosome I using CRISPR/Cas9 gene editing technology to monitor *nosr-1* expression (58). Actinomycin D treatment significantly increased the expression of *nosr-1_P_::NLS::GFP* (Figs. 6B, S12A). Knocking down *exos-9* by RNAi or mutations in exosome genes significantly increased the expression of *nosr-1_P_::NLS::GFP* (Figs. 6C-D, S12B-C). We measured the mRNA levels of endogenous *nosr-1* mRNA by quantitative real-time PCR and identified an increase in the mRNA levels of *nosr-1* (Figs. 6E-F). Similarly, knocking down rRNA biogenesis and ribosome assembly factors, the mutation of which induced NoSB formation, increased the expression of *nosr-1_P_::NLS::GFP* as well (Figs. S12D-G).

Taken together, these results suggested that nucleolar stress could activate the expression of *nosr-1*.

### *nosr-1* may engage in a new nucleolar stress response pathway

We tested whether other known stress response pathways are involved in the formation of nucleolar stress-induced NoSB. *cep-1* is an ortholog of mammalian p53 in *C. elegans* and has an essential role in nucleolar stress (34). SKN-1 senses nucleolar stress to activate xenobiotic detoxification genes (35). PHA-4 is a sensor of nucleolar stress and transactivates the expression of lipogenic genes (36); DAF-16 is a FOXO transcription factor involved in lifespan regulation as well as in oxidative stress and heat stress resistance (60, 61). However, none of these genes are required for *exos-10(ust242)* mutation-induced NoSB formation (Figs. S13A-B). Therefore, we speculated that *nosr-1* may engage in a new nucleolar stress response pathway.

### Transcriptome analysis identified *numr-1* as the NOSR-1 target

We performed mRNA-seq to explore the NOSR-1-targeted genes involved in NoSB formation. We first compared transcriptomes of wild-type and *nosr-1*(-) mutant animals under different stress conditions (Figs. 7A, S14A-C) and identified 11 genes that were consistently upregulated by nucleolar stress (Fig. 7B), which depend on the presence of functional NOSR-1 (Figs. 7C-E)

**Figure 7.**
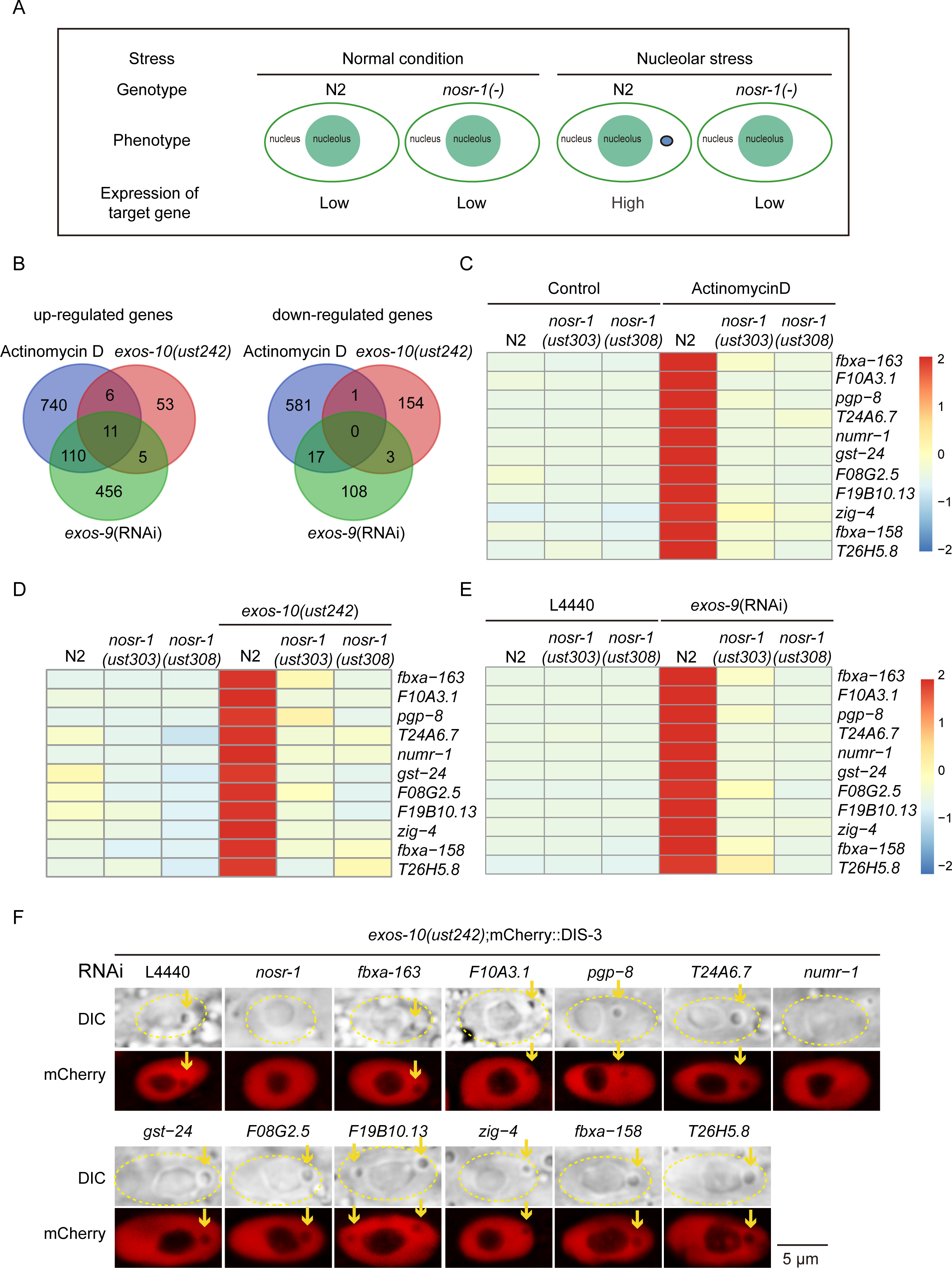
Transcriptome analysis identified NOSR-1 targets. (A) Schematic of the animal with or without nucleolar stress. (B) Venn diagrams revealing the overlapping target genes that are upregulated or downregulated under three nucleolar stress conditions. (C-E) Heatmap of the standardized fragments per kilobase of transcript per million mapped reads (FPKM) for the 11 coupregulated target genes in the indicated animals. The expression levels are indicated by the color bar. (F) DIC and fluorescence microscopy images of hypodermal nuclei in *exos-10(ust242)*;mCherry::dis-3 worms after knockdown of the indicated genes by RNAi.

We then tested whether nucleolar stress-induced NoSB depends on the 11 genes. We performed feeding RNAi experiments and knocked down the 11 genes in mCherry::DIS-3;*exos-10(ust242)* animals (Fig. 7F). RNAi knockdown of *numr-1* results in the prohibition of NoSB formation, suggesting that nucleolar stress-induced NoSB requires the presence of NUMR-1.

### Nucleolar stress induced the accumulation of NUMR-1 in NoSB

NUMR-1 (nuclear localized metal responsive) was previously identified as an SR-like protein that promotes cadmium tolerance (Fig. 8A) (62–64). NUMR-1 contains an RNA-recognition motif (RRM), serine- and arginine-rich regions that are common in SR proteins, and a C-terminal tail rich in histidine and glycine, suggesting that NUMR-1 might function in RNA metabolism (65–67). Interestingly, the SR family of proteins is strongly associated with liquid‒liquid phase separation (LLPS) and is enriched in nuclear bodies, especially in nuclear speckles (68). These features suggest that NUMR-1 might function in RNA metabolism and promote the formation of various biomolecular condensates.

**Figure 8.**
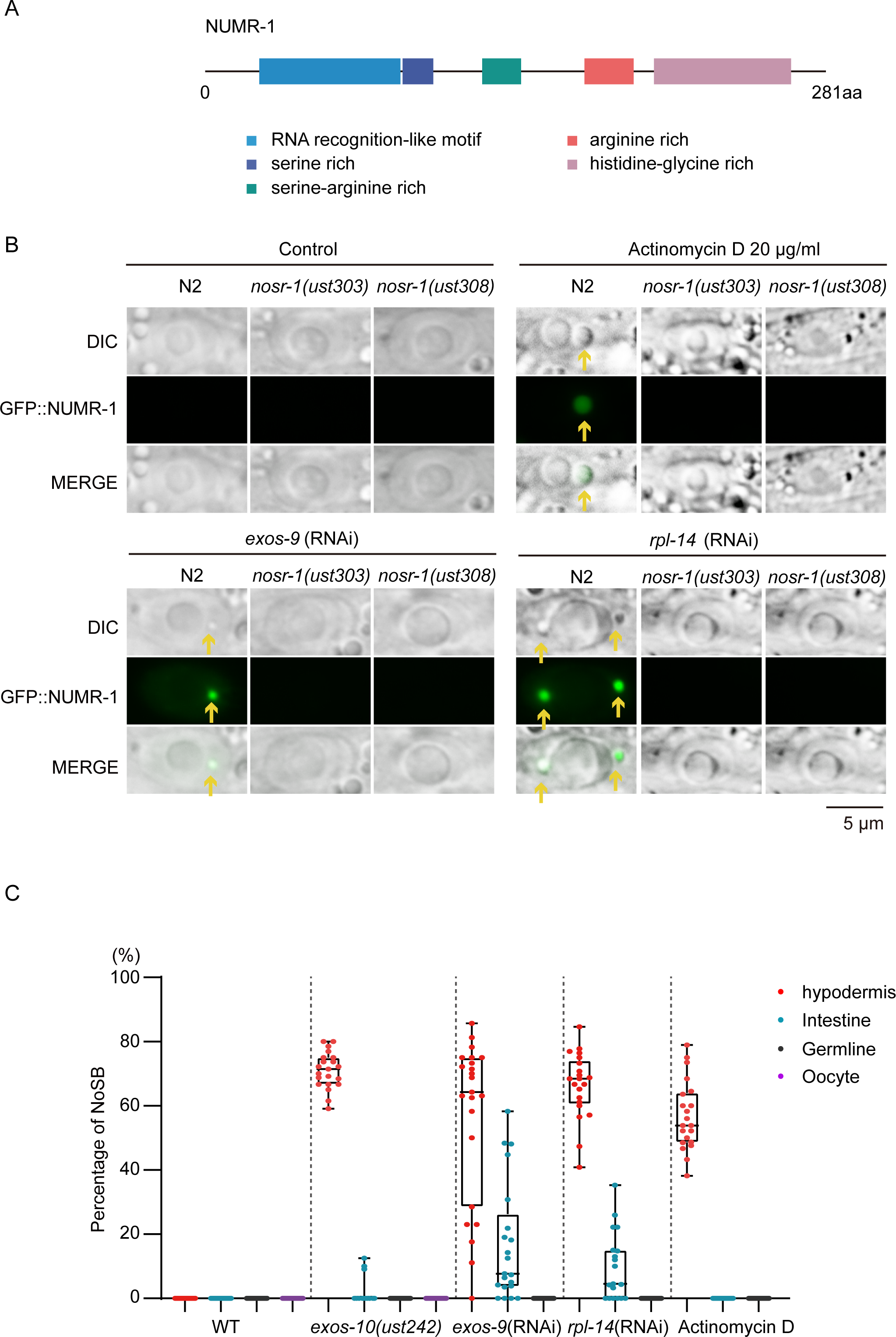
NUMR-1 is a component of NoSB. (A) Schematic of the *numr-1* protein structure. (B) DIC and fluorescence microscopy images of hypodermal nuclei after actinomycin D treatment and RNAi knockdown of *exos-9* and *rpl-14* in the indicated animals. (C) Quantification of NoSB in indicated tissues with indicated treatment. Every dot indicates the percentage of hypodermis cells containing NoSB in each worm. N > 20 animals, see also Figs. S16A-D.

To study the function of NUMR-1 in NoSB formation, we constructed a 3xFLAG-GFP–tagged NUMR-1 transgene (abbreviated GFP::NUMR-1) using CRISPR/Cas9-directed in situ gene editing technology. In wild-type animals cultured under normal laboratory conditions, we did not observe explicit GFP::NUMR-1 expression (Figs. 8B, S15A). Strikingly, upon nucleolar stress by the inhibition of rRNA transcription using actinomycin D or RNAi knockdown of *exos-9* or *rpl-14*, GFP::NUMR-1 expression was activated, and the protein accumulated in distinct nuclear foci under fluorescence microscopy (Figs. 8B, S15A). DIC microscopy indicated that NUMR-1 foci colocalized with NoSB (Fig. 8B). As expected, the *nosr-1* mutation inhibited GFP::NUMR-1 expression (Figs. 8B, S15A).

We also quantified the percentage of NoSB formation in different tissues (Figs. 8C and S16A-D). NoSB was mostly found in the hypodermal and intestinal cells, but rarely noticeable in germline and oocyte cells upon nucleolar stress.

To test whether NUMR-1 alone is sufficient to direct NoSB formation, we generated a *fib-1p::NUMR-1::GFP::fib-1utr* transgene to constitutively express NUMR-1 in all tissues. The ectopically expressed *fib-1p::*NUMR-1::GFP evenly diffused in the nucleoplasm without condensation into NoSB in the absence of nucleolar stress (Figs. S17A-B). The nucleolar stress treatment via Actinomycin D or *exos-9* RNAi induced the NoSB accumulation of *fib-1p::*NUMR-1::GFP, similar to the endogenous *numr-1* promoter-driven NUMR-1::GFP. Interestingly, this constitutively expressed *fib-1p::*NUMR-1::GFP did not rescue *nosr-1* mutation and failed to direct NoSB formation upon nucleolar stress. In the *nosr-1*(*-*)*; fib-1p::NUMR-1::GFP::fib-1UTR* animal, we did not observe noticeable condensation of NUMR-1::GFP and formation of NoSB. We speculated that there are additional NOSR-1-dependent factors besides NUMR-1 to synergistically mediate NoSB formation, therefore the constant expression of NUMR-1 only is not sufficient to rescue all *nosr-1*(*-*) defects. The identification of the components, both RNAs and proteins, in NoSB will greatly help to clarify the function and regulation of NoSB.

Together, these data suggest that NUMR-1 is a downstream target of NOSR-1 and is a crucial component of NoSB.

### NoSB may not obviously contain liquid-liquid phase separation (LLPS) properties

To test whether NUMR-1 can undergo phase separation, we first visualized the fine structure of NoSB by collecting Z-stack images of GFP::NUMR-1 in NoSB (Fig. 9A). GFP::NUMR-1 revealed even distribution throughout NoSB, and did not display noticeable subcompartments.

**Figure 9.**
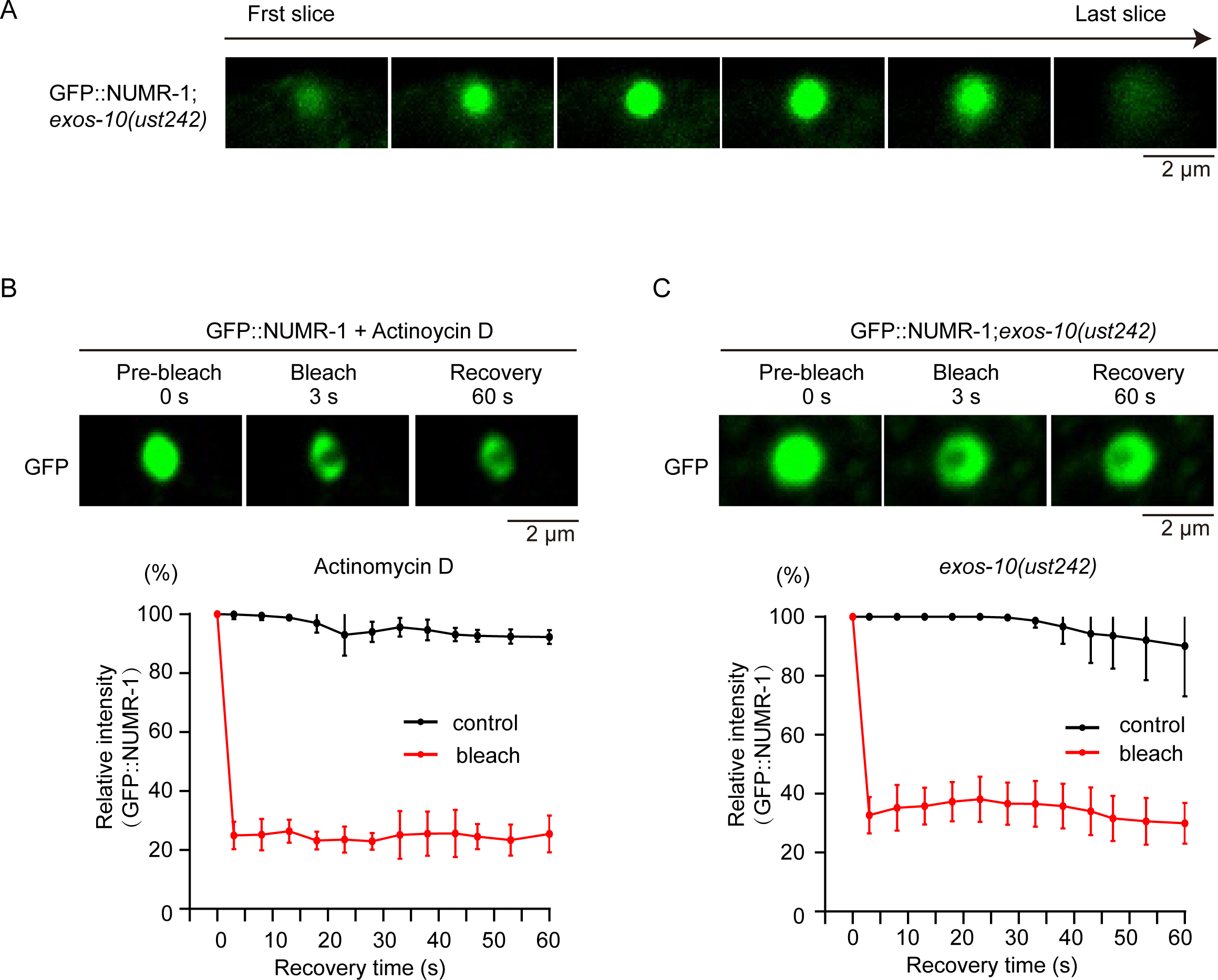
NoSB did not reveal obvious liquid-liquid phase separation property. (A) Z-stack images of the indicated animals with a step size of 0.5 μm. (B-C) (Top) FRAP assay of GFP::NUMR-1 transgene in indicated regions after Actinomycin D treatment or in *exos-10(ust242)* mutants. (Bottom) Quantification of FRAP data. Mean ± SD, n = 3.

Then, we induced NoSB formation via Actinomycin D treatment or *exos-10* depletion and performed fluorescence recovery after photobleaching (FRAP) assays (Figs. 9B-C). However, we failed to observe the recovery of GFP::NUMR-1 foci after photobleaching. These data suggested that NoSB may not obviously contain liquid-liquid phase separation (LLPS) properties.

To explore the recovery dynamics of NoSB from nucleolar stress, we treated worms with actinomycin D for 48 hours, then transferred the stressed worms to normal growth media without actinomycin D, and quantified the percentage of NoSB thereafter (Figs. S18A-C). As expected, the percentage of NoSB as well as the expression of GFP::NUMR-1 gradually decreased upon the removal of actinomycin D (Figs. S18B-C), suggesting that constant nucleolar stress is required for the maintenance of NoSB condensation.

Using NUMR-1::GFP as the marker, we tested again whether other kinds of environmental conditions besides nucleolar stress could induce the formation of NoSB. However, growing animals at 15°C or 25°C, cold shock at 4°C or heat shock at 37°C failed to induce the formation of NoSB (Figs. S19A-B, also see S3E-G). Yet occasionally, we observed one or two small GFP::NUMR-1 foci in a few animals at normal growing condition as well as under certain environmental stresses (Figs. S19A-B), which were hardly detectable by DIC.

### Cadmium stress induced NUMR-1 expression and *nosr-1*-independent NoSB formation

NUMR-1 (nuclear localized metal responsive) was previously identified as an SR-like protein that promotes cadmium tolerance (62–64). Cadmium was shown to increase *numr-1/-2* mRNA expression in pharyngeal and intestinal cells (69). Cadmium arrests larval development and decreases body size, and these effects were exacerbated by *numr-1/-2* RNAi (62).We then tested whether cadmium exposure could translocate NUMR-1 to NoSB.

As expected, the treatment with cadmium increased the expression of GFP::NUMR-1 (Fig. S20A). In addition, cadmium induced the aggregation of GFP::NUMR-1 in the nucleus in intestine cells, but not in hypodermis (Figs. 10A-B). Interestingly, the cadmium-induced NUMR-1 expression and condensation did not dependent on *nosr-1*. In *nosr-1* mutants, cadmium could still stimulate NUMR-1 expression and aggregation (Figs. 10A-B, S20A), suggesting a *nosr-1*-independent pathway is involved in the formation of cadmium-induced NUMR-1 condensation.

**Figure 10.**
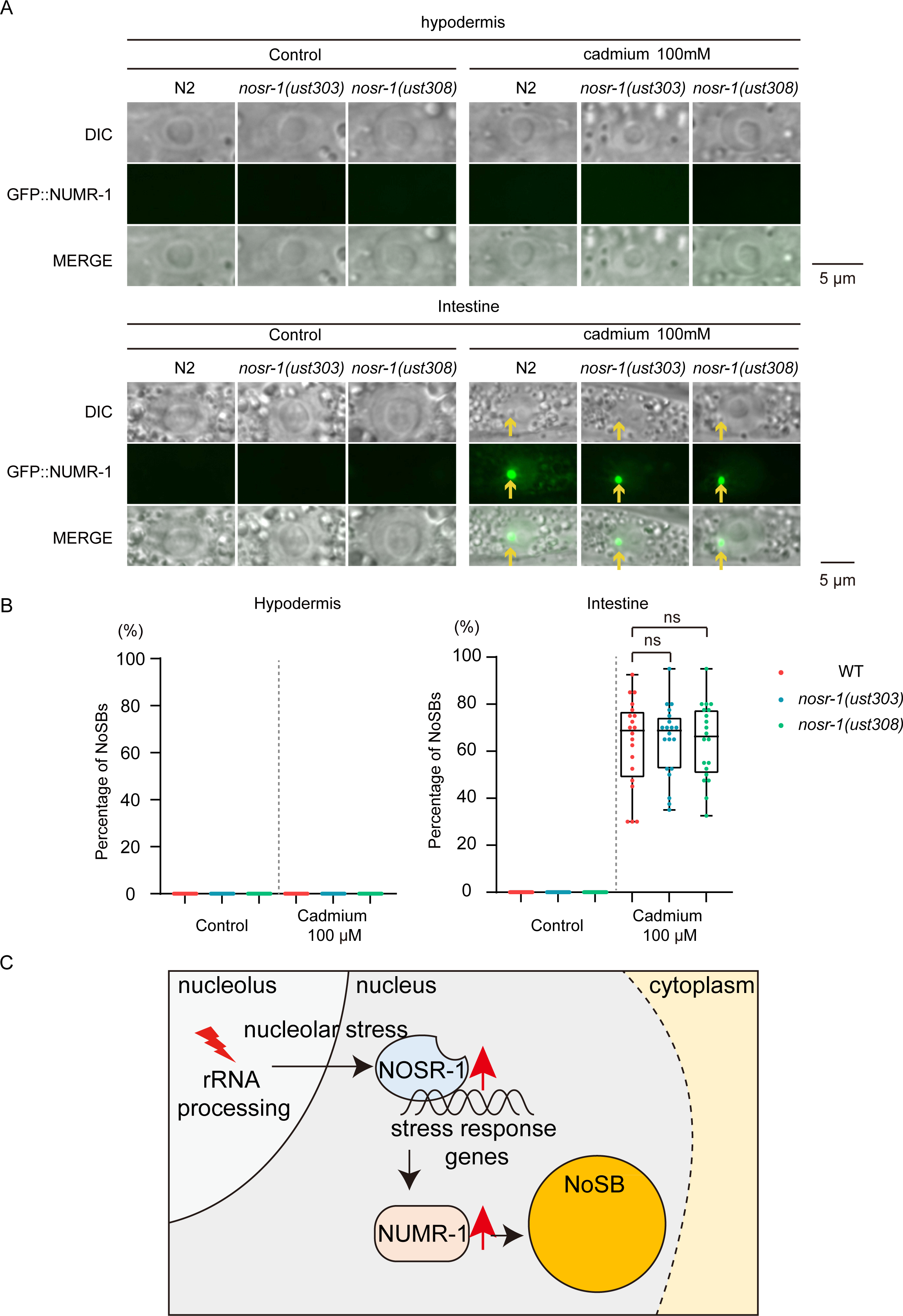
Cadmium stress induced NoSB formation in a *nosr-1*-independent manner. (A) DIC and fluorescence microscopy images of hypodermal and intestine nuclei after cadmium treatment in the indicated animals. (B) Quantification of NoSB in hypodermic cells and intestine cells from (A). Every dot indicates the percentage of hypodermis or intestine cells containing NoSB in each worm. n>19 animals. (C) A working model of NoSB formation under nucleolar stress in *C. elegans*.

## Discussion

Environmental stresses not only alter gene expression but also trigger structural changes in cells. However, how distinct subcellular structures respond to stresses remains largely mysterious. Here, we showed that nucleolar stress induced the formation of a new subnuclear organelle, nucleolar stress body (NoSB), which is regulated by a new bZIP transcription factor, NOSR-1. Further experiments identified that NUMR-1 is the critical component of NoSB, is induced by NOSR-1 and is per se required for NoSB formation (Fig. 10C), suggesting that NUMR-1 may be involved in building up the scaffold and recruiting NoSB components.

### Nucleolar stress induced the formation of a nucleolar stress body via the NOSR-1/NUMR-1 axis

In mammals, the tumor suppressor protein p53/CEP-1 is considered an essential factor in monitoring nucleolar stress and the integrity of ribosome biogenesis (20). Additional pathways have also been identified to respond to nucleolar stresses in the absence of functional p53 (67, 68). The ubiquitin E3 ligase MDM2 negatively regulates p53 by marking it for ubiquitin-mediated proteasomal degradation. In plant cells, nucleolar stress activates the expression of ANAC082 by releasing ribosome stalling at the upstream open reading frame (uORF) (31). These multiple layers of regulation enable transcription factors to execute cellular responses to specific nucleolar stress conditions. Here, we found that nucleolar stress activates the expression of NOSR-1 and identified a novel NOSR-1/NUMR-1-dependent pathway that links nucleolar stress with the formation of NoSB. However, how the NOSR-1/NUMR-1 axis senses perturbed ribosome biogenesis and nucleolar disorders is unclear.

NUMR-1 is upregulated by NOSR-1 and is required for NoSB formation upon nucleolar stress, suggesting that NUMR-1 functions downstream of NOSR-1. We also identified a diverse set of mRNAs that are upregulated by NOSR-1 but do not participate in NoSB formation, indicating that NOSR-1 may be involved in a variety of additional cellular processes. Further experiments, for example, chromatin immunoprecipitation followed by deep sequencing (ChIP-seq), are required to identify the direct targets of NOSR-1.

### NUMR-1 and the nucleolar stress body

Membraneless organelles (MLOs) are condensates thought to form through liquid‒ liquid phase separation (LLPS) via various mechanisms, for example, protein/RNA interactions (2, 70, 71). The key component of cytoplasmic stress granules (SGs), G3BP1, has a C-terminal RNA-binding domain (RBD), which functions as a molecular switch to trigger RNA-dependent LLPS in response to a rise in intracellular free RNA concentrations (72).

NUMR-1 encodes RRM and serine- and arginine-rich regions that are found in canonical SR proteins (62). Many SR proteins have reported functions in protein/RNA nucleation. For example, the lncRNA metastasis-associated lung adenocarcinoma transcript 1 (MALAT1) localizes to nuclear speckles where it interacts with SR proteins (73). SRSF1 and SRSF9 assemble nSBs when cells are exposed to high temperatures (74). Some SR proteins (SRSF1, SRSF2, SRSF3, SRSF7, and SRSF10) are recruited to SGs along with nontranslated mRNAs (75). Whether and how NUMR-1 uses a similar mechanism to promote NoSB formation requires further investigation.

The components of NoSB are very intriguing. Till now, we found that (1) treating worms with Actinomycin D or inactivation of RNA exosome did not directly translocate NUCL-1 or NUCL-1(ΔRGG) into NoSB. However, a further treatment with NaN_3_ after Actinomycin D or inactivation of RNA exosome could induce an accumulation of NUCL-1 or NUCL-1(ΔRGG) to NoSB. (2) A SR-like protein NUMR-1 is specifically enriched in NoSB after nucleolar stress via Actinomycin D, inactivation of RNA exosome, or cadmium treatment.

NoSB and GFP::NUMR-1 are rarely detectable in steady-state cells; however, their expression drastically increases in the hypodermal cells and intestinal cells upon nucleolar stress. The NoSB also disassembles following stress removal. Therefore, we speculated that NUMR-1 might recruit stress-induced proteins and RNAs to NoSB upon nucleolar stress. How NoSB interact with stress responsive proteins and RNAs is unknown. Further investigation to identify the protein and RNA components in NoSB is crucial to elucidate the functions of NoSB and the underlying mechanisms.

NaN_3_ can quickly immobilize nematodes, which makes it an ideal tool to paralyze worms and is widely used in imaging experiments (76, 77). Treating *C. elegans* with NaN_3_ induces hypoxia and eventually leads to cell death (78). For all the proteins tested in the work, including markers for nucleolus and other subnuclear organelles, we only observed that NUCL-1 and NUCL-1(ΔRGG) aggregated to NoSB after the treatment of NaN_3_ following the nucleolar stress. yet NUCL-1 is not required for NoSB formation. The reason is not clear yet. Nucleolin likely functions in rRNA transcription and processing. In *C. elegans*, NUCL-1 located in GC region and is required for nucleolar vacuole formation (37). Whether and how NaN_3_-induced hypoxia or cell death might change the biochemical properties or binding partners of NUCL-1 require further investigation.

### The function of NoSB

Membrane organelles were thought to increase the efficiency of cellular processes by increasing the local concentration of proteins, RNAs, and even committed genes in a dedicated microenvironment (79). Processes and pathways that are known to be regulated by membrane organelles include many important facets of cellular functions, including genome organization and repair, RNA splicing, gene expression, and ribosome production (80–82). Impairment in the assembly of membrane organelles may be harmful for organisms. For example, the depletion of the SG core component G3BP1 in neurons leads to an increase in intracellular calcium and calcium release and neuronal dysfunction (83). Upon the depletion of Cajal bodies, immortal and highly proliferative aneuploid cancer cells display genome-wide RNA splicing defects, which are detrimental to long-term cell viability (84).

NUMR-1 was previously identified as an SR-like protein that promotes cadmium tolerance (62–64). Cadmium arrests larval development and decreases body size, and these effects were exacerbated by *numr-1/2* RNAi (62). Cadmium was shown to increase *numr-1/2* mRNA expression (69). Therefore, we speculated that NUMR-1/2 and NoSB may involve in protecting animals from adverse environment conditions. The identification of the components, both RNAs and proteins, in NoSB will greatly help to clarify the function and regulation of NoSB.

## Materials and methods

### Strains

Bristol strain N2 was used as the standard wild-type strain. All strains were grown at 20°C unless otherwise specified. The strains used in this study are listed in Supplementary Table S2.

### Construction of transgenic strains

For ectopic transgene expression of *nosr-1_p_::NLS::GFP*, the NOSR-1 promoter and UTR were amplified from N2 genomic DNA. The EGL-3 NLS sequence was amplified from N2 genomic DNA. A GFP::3xFLAG sequence was PCR amplified from SHG326 genomic DNA. The vector fragment was PCR amplified from the plasmid pSG274. These fragments were joined together by Gibson assembly to form the repair plasmid with the ClonExpress MultiS One Step Cloning Kit (Vazyme Biotech, Nanjing, China, Cat. No. C113-01/02). The transgene was integrated into *C.* elegans chromosome I via a modified counterselection (cs)-CRISPR method. The sequences of the primers are listed in Supplementary Table S3.

For in situ knock-in transgenes, the 3xFLAG::GFP sequence was PCR amplified from YY178 genomic DNA. The GFP::3xFLAG sequence was PCR amplified from SHG326 genomic DNA. The tagRFP coding sequence was PCR amplified from YY1446 genomic DNA. Homologous left and right arms (1.5 kb) were PCR amplified from N2 genomic DNA. The backbone was PCR amplified from the plasmid pCFJ151. All these fragments were joined together by Gibson assembly to form the repair plasmid with the ClonExpress MultiS One Step Cloning Kit. The plasmid was coinjected into N2 animals with three sgRNA expression vectors, 5 ng/µl pCFJ90, and the Cas9 II-expressing plasmid. The sequences of the primers are listed in Supplementary Table S4.

### Construction of deletion mutants

For gene deletions, triple sgRNA-guided chromosome deletion was conducted as previously described (58). To construct sgRNA expression vectors, the 20 bp *unc-119* sgRNA guide sequence in the pU6::unc-119 sgRNA(F+E) vector was replaced with different sgRNA guide sequences. Addgene plasmid #47549 was used to express Cas9 II protein. Plasmid mixtures containing 30 ng/µl of each of the three or four sgRNA expression vectors, 50 ng/µl Cas9 II-expressing plasmid, and 5 ng/µl pCFJ90 were coinjected into N2 animals. Deletion mutants were screened by PCR amplification and confirmed by sequencing. The sgRNA sequences are listed in Supplementary Table S5.

### Imaging and quantification

Images were collected on a Leica DM4 B microscope. The nucleolar area and NoSB area were quantified manually with the freehand tool using Fiji software.

### Actinomycin D treatment

Actinomycin D (MedChemExpress no. HY-17559, CAS: 50-76–0) was prepared at 20 mg/ml in DMSO as a stock solution. The actinomycin D stock solution was diluted to 5 to 30 μg/ml with concentrated OP50. NGM plates were prepared and placed at room temperature overnight before use. Synchronized L1 worms were placed onto the seeded plates and grown for 48 hours before imaging.

### RNAi

RNAi experiments were performed at 20°C by placing synchronized embryos on feeding plates as previously described (85). HT115 bacteria expressing the empty vector L4440 (a gift from A. Fire) were used as controls. Bacterial clones expressing dsRNAs were obtained from the Ahringer RNAi library and sequenced to verify their identity. Some bacterial clones were constructed in this work, which are listed in Supplementary Table S1. Images were collected using a Leica DM4 B microscope.

### Forward genetic screening

Forward genetic screening was conducted as previously described (86). To identify factors prohibiting the formation of NoSB, we chemically mutagenized the *mCherry::DIS-3* strain by ethyl methanesulfonate (EMS), followed by a clonal screen. The F2 progeny worms were visualized under a fluorescence microscope at the L4 stage. Two mutants that exhibited NoSB formation were isolated from two thousand haploid genomes. *exso-10 (ust242)* and *exos-4.2 (ust243)* were identified by genome resequencing.

To identify factors required for the formation of NoSB, we chemically mutagenized the *exso-10(ust242)*;*mCherry::DIS-3* strain by ethyl methanesulfonate (EMS), followed by a clonal screen. The F2 progeny worms were visualized under a fluorescence microscope at the L4 stage. One mutant that disrupted NoSB formation was isolated from two thousand haploid genomes. *nosr-1(ust303)* was identified by genome resequencing.

### Postfix Oil Red O staining and bright field imaging

Postfix Oil Red O staining and bright field imaging were conducted as described previously (87). Briefly, L4 stage animals were fixed in 1% paraformaldehyde/PBS for 30 min with rocking. Samples were then frozen on dry ice/ethanol and thawed with running tap water three times. Samples were washed three times with 1X PBS, dehydrated in 60% isopropanol for 2 min, and stained with 0.5 ml 60% Oil-Red-O (#A6000395-50G, BBI) working solution for 30 min with rocking. Oil Red O stock solution was dissolved in isopropanol at a concentration of 0.5 g/100 ml and equilibrated for several days. The Oil-Red-O working solution was prepared fresh by mixing 60% volume of stock with 40% volume of water. The mix was equilibrated for 10 min and then filtered with a 0.22 μm syringe filter. Stained samples were then washed three times with 1X PBS, rehydrated in 1X PBS, and mounted in 1X PBS onto agarose-padded slides for imaging.

### RNA isolation

Synchronized L4 worms were incubated with TRIzol reagent followed by 5 quick liquid nitrogen freeze‒thaw cycles. RNA was precipitated by isopropanol followed by DNaseI digestion (Qiagen).

### mRNA deep-sequencing

The Illumina-generated raw reads were first filtered to remove adaptors, low-quality tags, and contaminants to obtain clean reads at Novogene. The clean reads were mapped to the reference genome of WBcel235 via HISAT2 software (version 2.1.0) (88). Differential expression analysis was performed using custom R scripts. A 2-fold-change cutoff was applied when filtering for differentially expressed genes. All plots were drawn using custom R scripts.

### qRT–PCR

cDNAs were generated from RNA using HiScript III RT SuperMix for qPCR (Vazyme), which includes a random primer/oligo(dT)20VN primer mix for reverse transcription. qPCR was performed using a MyIQ2 real-time PCR system (Bio-Rad) with AceQ SYBR Green Master mix (Vazyme). The primers used in qRT‒PCR are listed in Supplementary Table S4.

### Brood Size

L4 hermaphrodites were singled onto NGM plates and transferred daily as adults until embryo production ceased, and the progeny numbers were scored.

### Lifespan assay

Lifespan assays were performed at 20°C as described previously (89). Basically, worm populations were synchronized by placing young adult worms on NGM plates for 4–6 hours and then removing them. The hatching day was counted as day one for all lifespan measurements. Worms were transferred every other day to new plates to eliminate confounding progeny. Animals were scored as alive or dead every day. Worms were scored as dead if they did not respond to repeated prods with a platinum pick. Worms were censored if they crawled off the plates or died from vulval bursting and bagging. For each lifespan assay, >50 worms were used.

### Hydrogen peroxide assay

Ten synchronized worms on day 1 of adulthood were transferred to each well, which contained 1 ml of worm S-basal buffer with various concentrations of H_2_O_2_ in a 12-well plate at 20°C. Four hours later, 100 μl of 1 mg/ml catalase (Sigma, C9322) was added to neutralize H_2_O_2_, and the mortality of worms was scored.

### Heat-shock assay

Ten synchronized hermaphrodites on day 1 of adulthood were transferred to new NGM plates and incubated at 35°C, and their mortality was checked every 1 hr until there were no living worms.

### Starvation assay

L4 worms were transferred to empty NGM plates without any bacterial lawn for 12 hours, and then the animals were imaged.

### Cadmium treatment

Synchronized L1 worms were fed with OP50 on NGM agar plates containing 0, 100 or 300 μM of cadmium chloride for 48h before imaging.

### Fluorescence recovery after photo bleaching (FRAP)

FRAP experiments were performed using a Zeiss LSM980 laser scanning confocal microscope at room temperature. Worms were anesthetized with 2 mM levamisole. A region of interest was bleached with 70% laser power for 3–4 s, and the fluorescence intensities in these regions were collected every 5 s and normalized to the initial intensity before bleaching. For analysis, image intensity was measured by Mean and further analyzed by GraphPad Prism 9.0 software.

### Statistics

The mean and standard deviation of the results are presented in bar graphs with error bars. All experiments were conducted with independent *C. elegans* animals for the indicated number (N) of times. Statistical analysis was performed with the two-tailed Student’s t test or unpaired Wilcoxon test as indicated.

### Data availability

All the high throughput data were deposited onto the Genome Sequence Archive in the National Genomics Data Center, China National Center for Bioinformation/Beijing Institute of Genomics, Chinese Academy of Sciences (GSA: CRA012745), which are publicly accessible at https://bigd.big.ac.cn/gsa/browse/CRA012745.

## Acknowledgments

We are grateful to the members of the Guang lab for their comments. We are grateful to the International *C. elegans* Gene Knockout Consortium and the National Bioresource Project for providing the strains. Some strains were provided by the CGC, which is funded by the NIH Office of Research Infrastructure Programs (P40 OD010440). This work was supported by grants from the National Key R&D Program of China (2022YFA1302700 and 2019YFA0802600) and the National Natural Science Foundation of China (32230016, 32270583, 32070619, 2023M733425 and 32300438), and the Strategic Priority Research Program of the Chinese Academy of Sciences (XDB39010600), the Research Funds of Center for Advanced Interdisciplinary Science and Biomedicine of IHM (QYPY20230021) and the Fundamental Research Funds for the Central Universities.

## Supplemental figure legends

**Figure S1.**
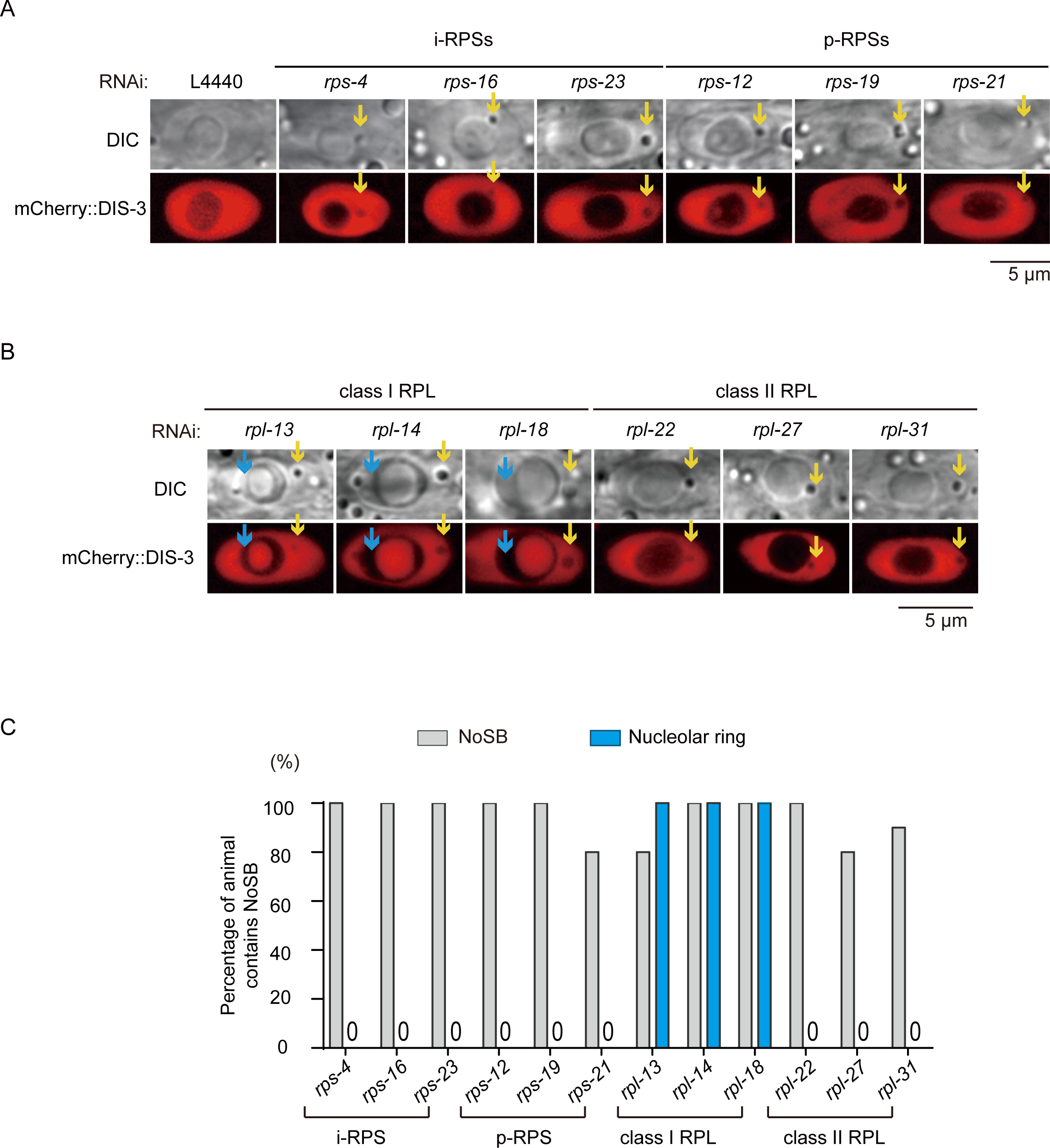
Candidate-based RNAi screening identified genes that suppress NoSB formation. (A-B) DIC and fluorescence microscopy images of nuclei after knockdown of the indicated genes by RNAi in the indicated animals. (C) Quantification of NoSB and ring-shaped nucleoli after RNAi knockdown of the indicated genes. The percentage of animals containing at least three NoSB structure were shown. n>19 animals.

**Figure S2.**
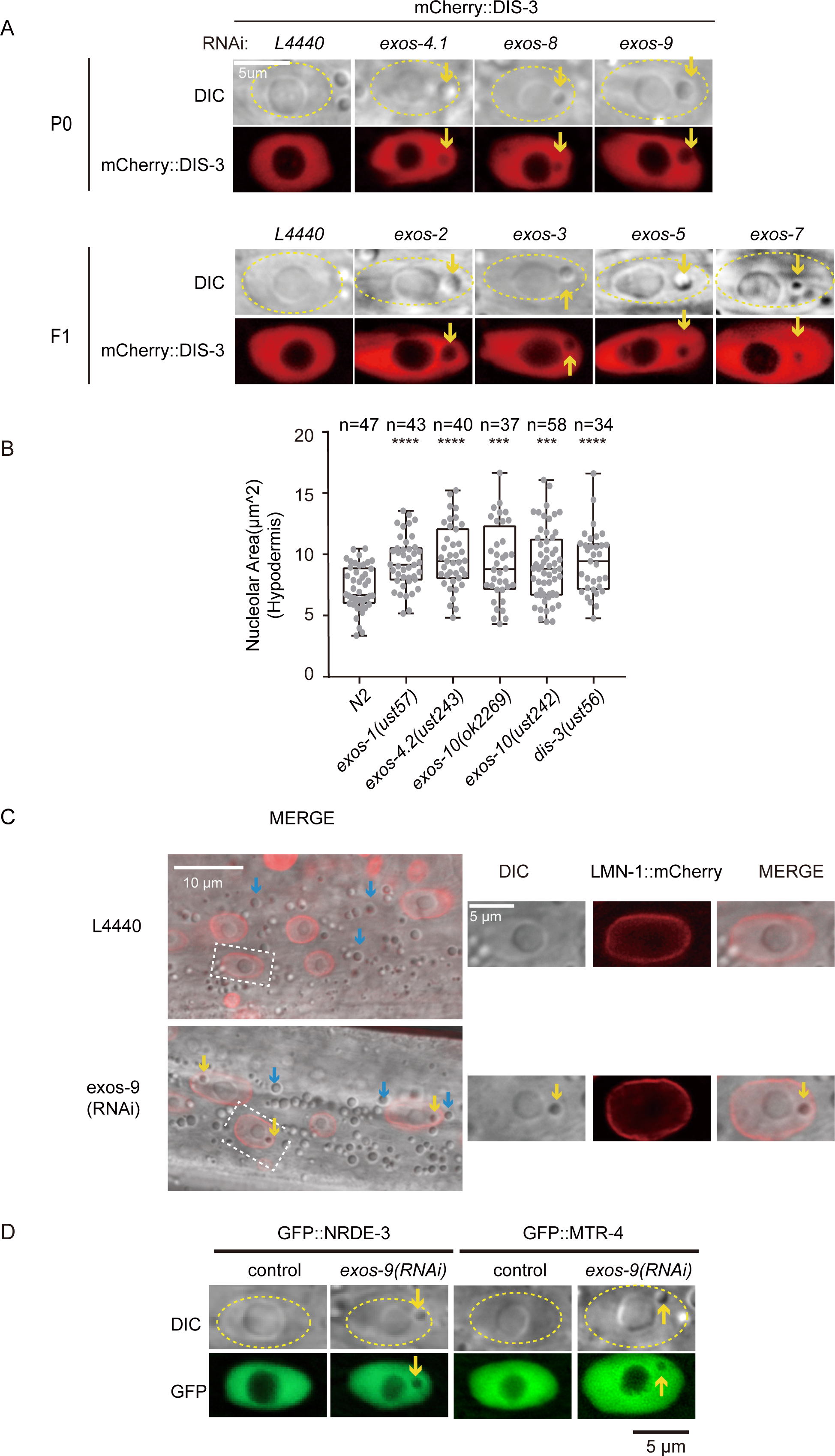
The depletion of exosome ribonucleases induced NoSB formation. (A) DIC and fluorescence microscopy images of nuclei after knockdown of the indicated genes by RNAi. (B) The size of the nucleoli in exosome mutants. n>30 nucleoli from n>19 animals. (C) DIC and fluorescence microscopy images of nuclei in LMN-1::mCherry transgenic animals after *exos-9 knockdown*. The blue arrows indicate fat droplets, and the yellow arrows indicate NoSB. (D) DIC and fluorescence microscopy images of nuclei in the indicated animals.

**Figure S3.**
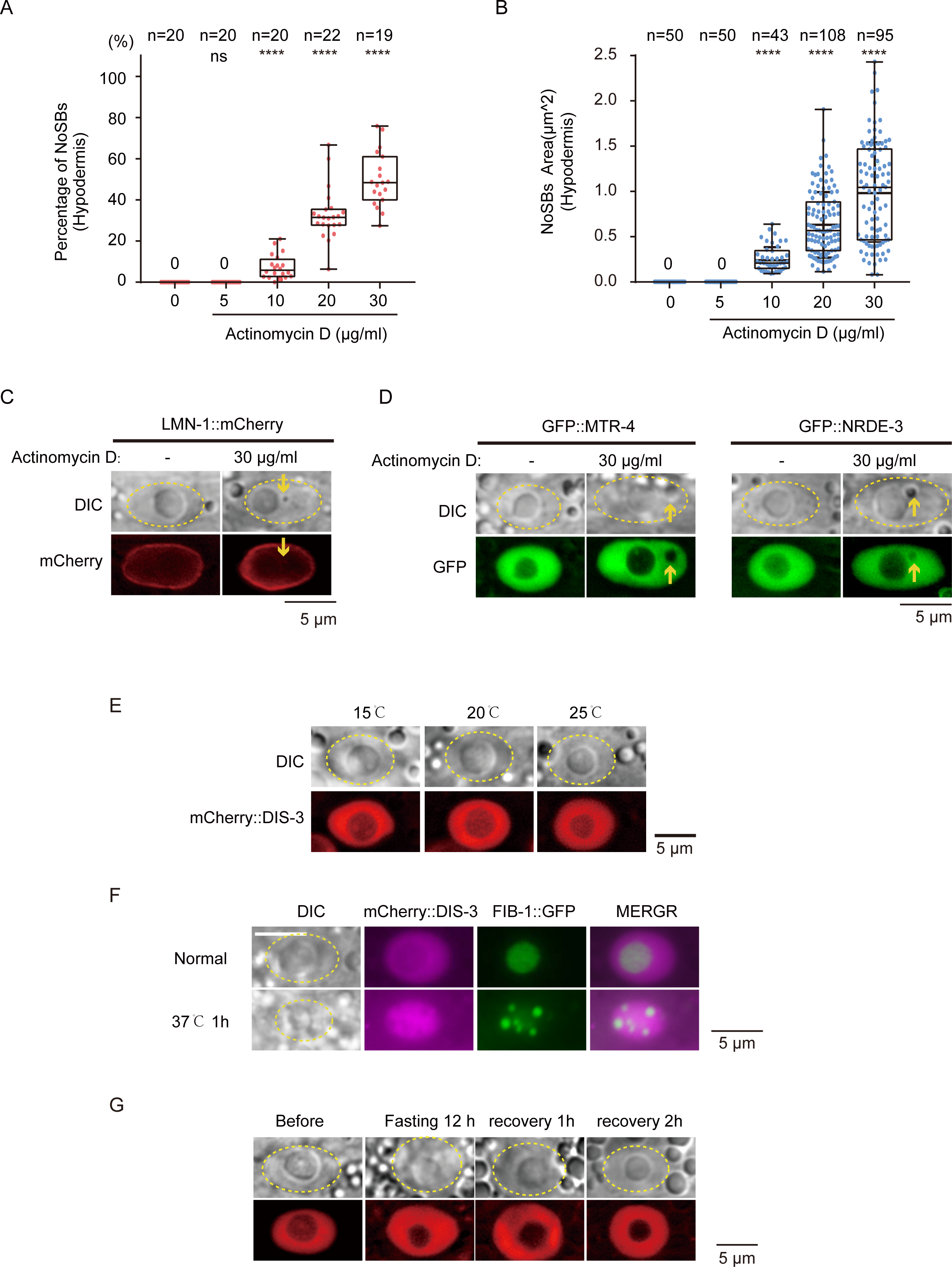
Temperature shifts could not induce the formation of NoSB. (A) Quantification of NoSB in hypodermic cells after actinomycin D treatment. n>18 animals. Every dot indicates the percentage of hypodermis cells containing NoSB in each worm. (B) The size of NoSB in hypodermic cells after actinomycin D treatment. n>40 NoSBs from n>18 animals. (C-D) DIC and fluorescence microscopy images of nuclei in the indicated animals after actinomycin D treatment. (E) DIC and fluorescence microscopy images of nuclei at the indicated temperatures. (F) DIC and fluorescence microscopy images show epidermal cells of the indicated L4 stage animals expressing mCherry::DIS-3 after 37°C heat shock for 1 hour. (G) DIC and fluorescence microscopy images of nuclei after starvation for 12 h.

**Figure S4.**
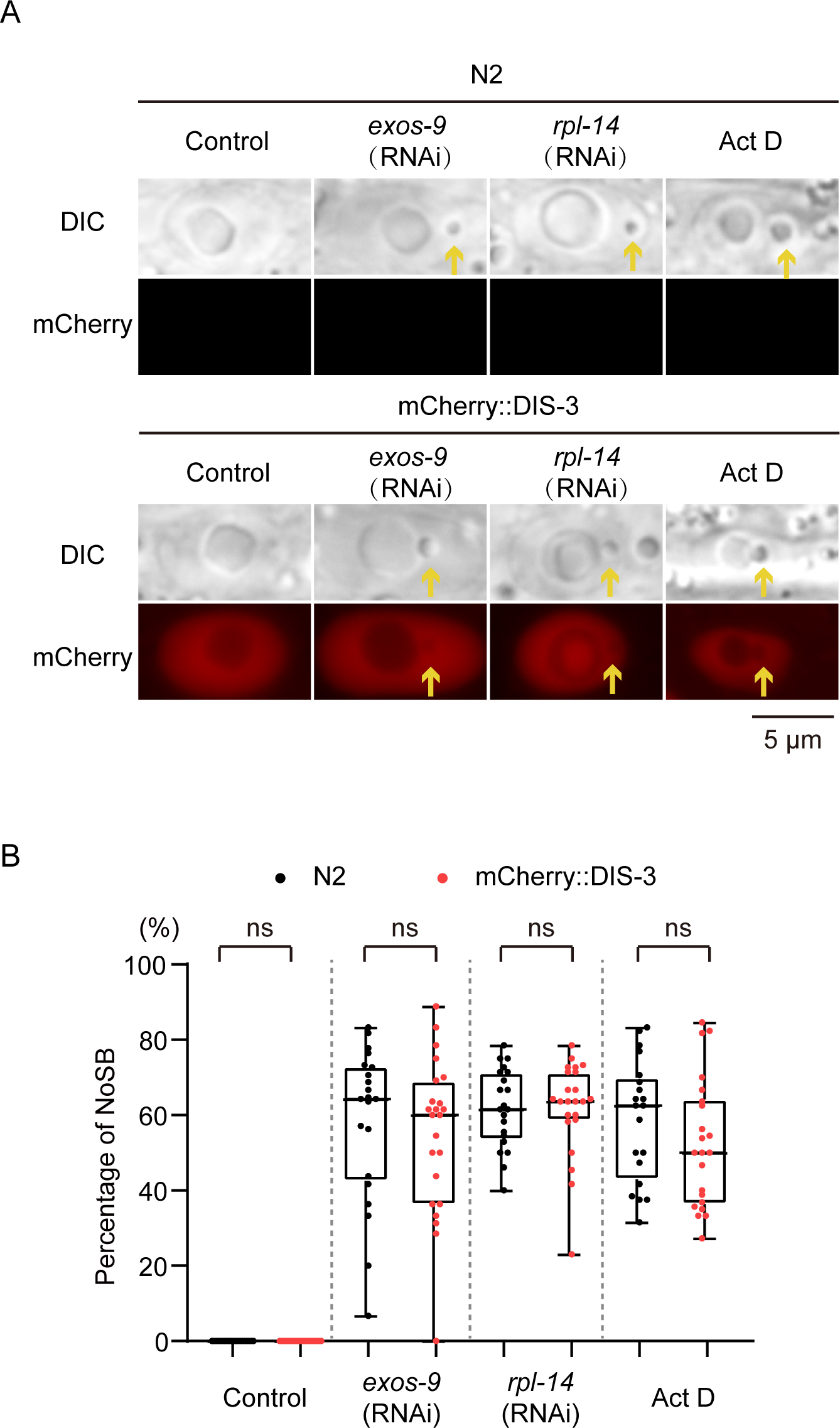
The ectopically expressed mCherry::DIS-3 does not affect NoSB formation. (A) DIC and fluorescence microscopy images of nuclei in hypodermal cells after knockdown of the indicated genes or Actinomycin D treatment in the indicated animals. (B) Quantification of the NoSB in indicated animals upon indicated treatment. N > 20 animals. Every dot indicates the percentage of hypodermis cells containing NoSB in each worm.

**Figure S5.**
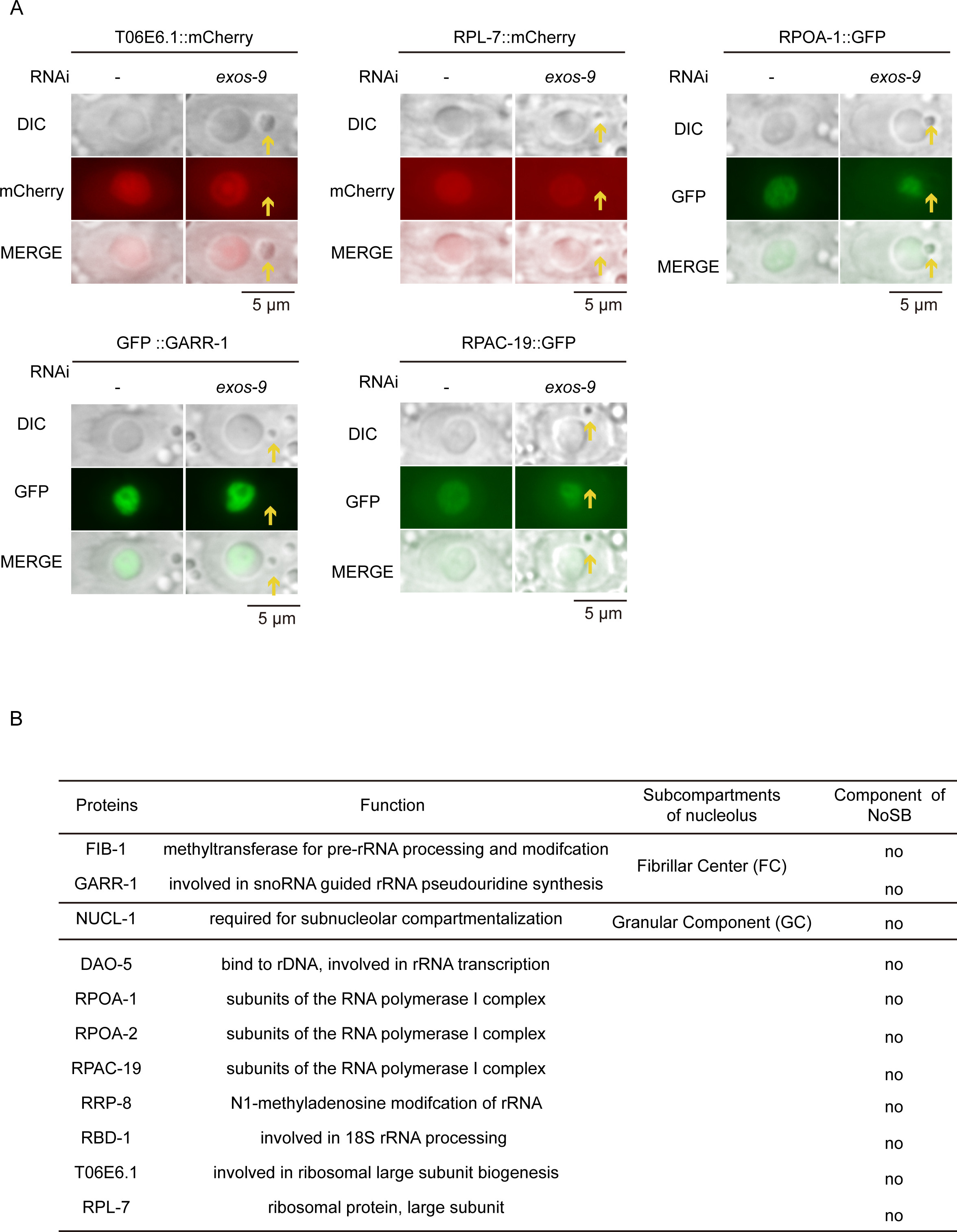
NoSB is not a portion of the nucleolus that blebbed off upon nucleolar stress. (A) DIC and fluorescent microscopy images of indicated transgenes after knocking down *exos-9*. (B) Summary of the localization of indicated proteins after *exos-9* RNAi.

**Figure S6.**
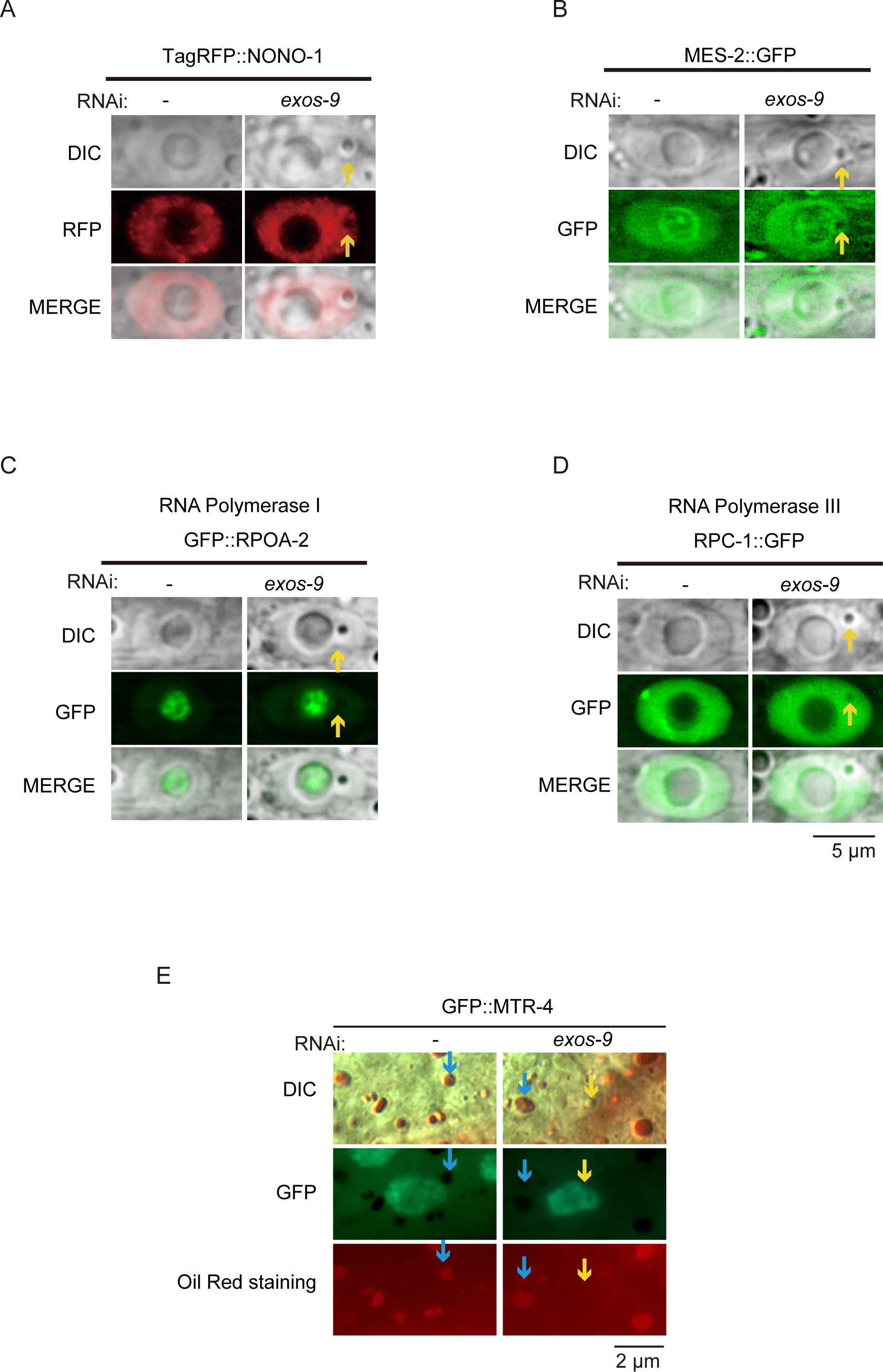
NoSB is likely a new subnuclear organelle. (A-D) DIC and fluorescence microscopy images of hypodermal nuclei with the indicated transgenes after *exos-9 knockdown*. (E) Oil Red O-stained lipid droplets in fixed GFP::MTR-4 transgenic worms with or without *exos-9* RNAi. The blue arrows indicate fat droplets, and the yellow arrows indicate NoSBs.

**Figure S7.**
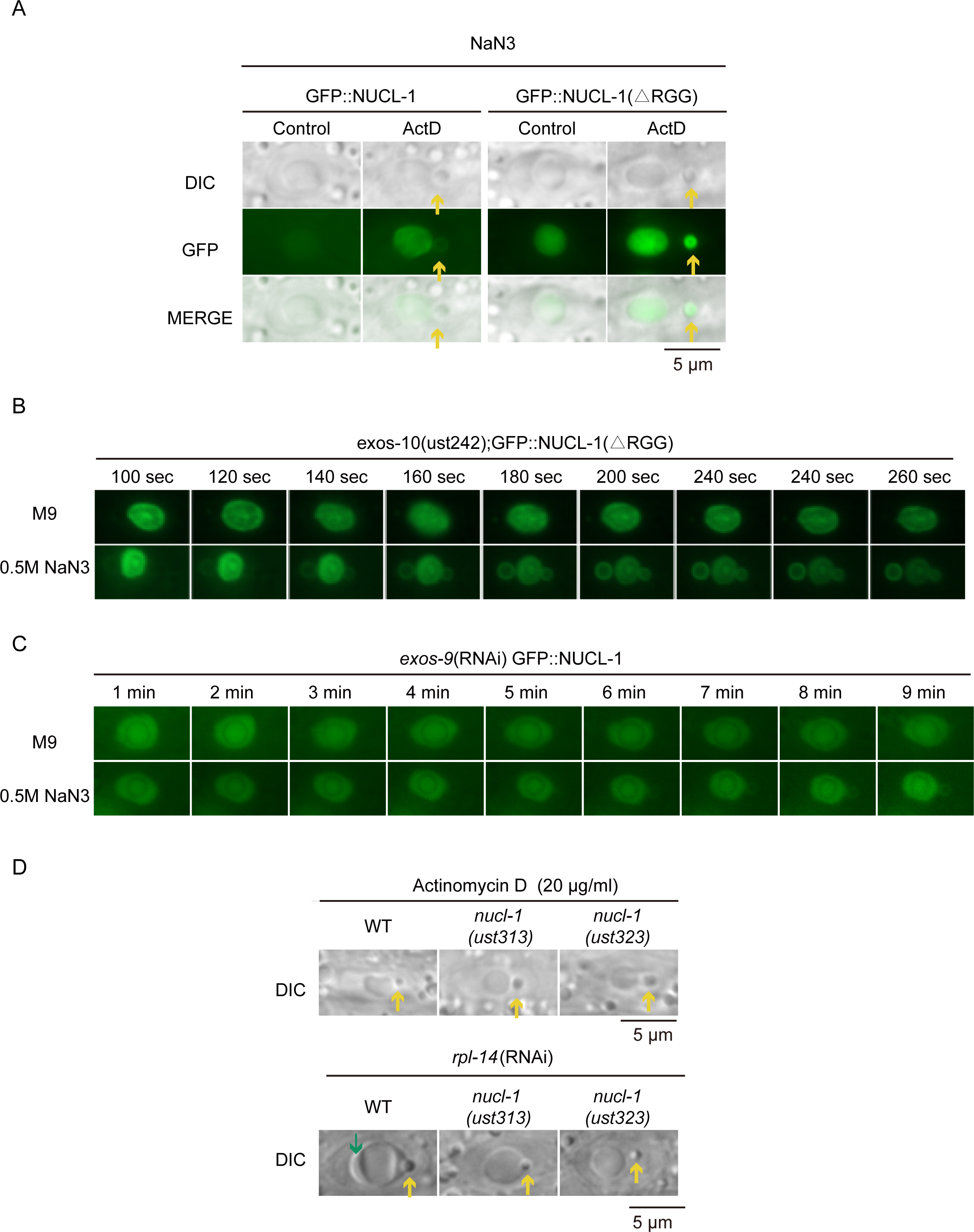
NaN_3_ and nucleolar stress co-treatment induced the translocation of NUCL-1 to NoSB. (A) DIC and fluorescence microscopy images of *C. elegans* nuclei after actinomycin D followed by NaN_3_ treatment. (B-C) Fluorescence microscopy images of *C. elegans* nuclei after the depletion of exosome followed by NaN_3_ treatment. (D) DIC images of *C. elegans* nuclei after actinomycin D or *rpl-14* RNAi in indicated animals.

**Figure S8.**
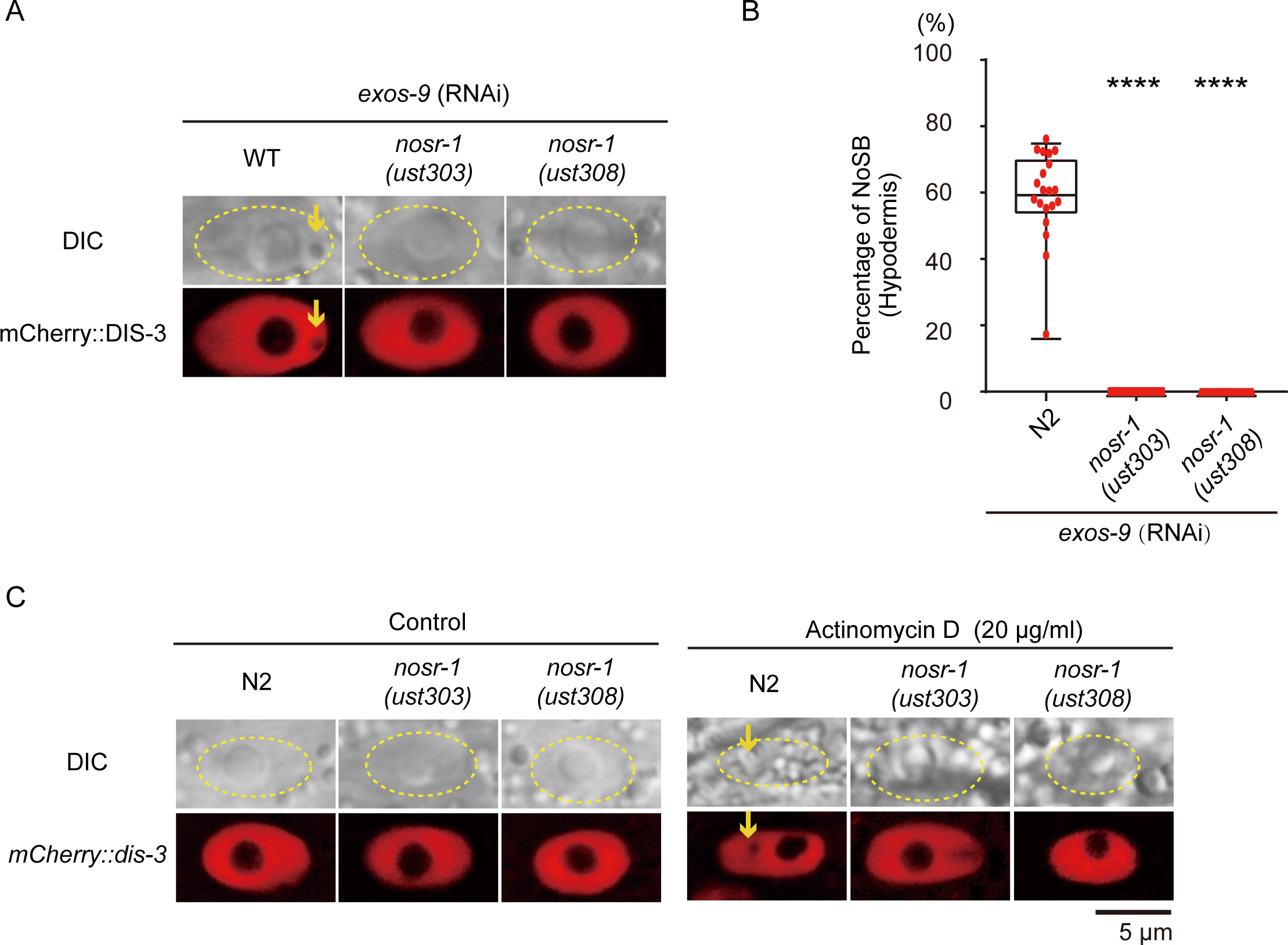
NOSR-1 is required for NoSB formation. (A) DIC and fluorescence microscopy images of *C. elegans* nuclei after knocking down *exos-9* by RNAi in *nosr-1* mutants. (B) Quantification of NoSB in hypodermic cells from (A). Every dot indicates the percentage of hypodermis cells containing NoSB in each worm. n>19 animals. (C) DIC and fluorescence microscopy images of nuclei after actinomycin D treatment in the indicated animals.

**Figure S9.**
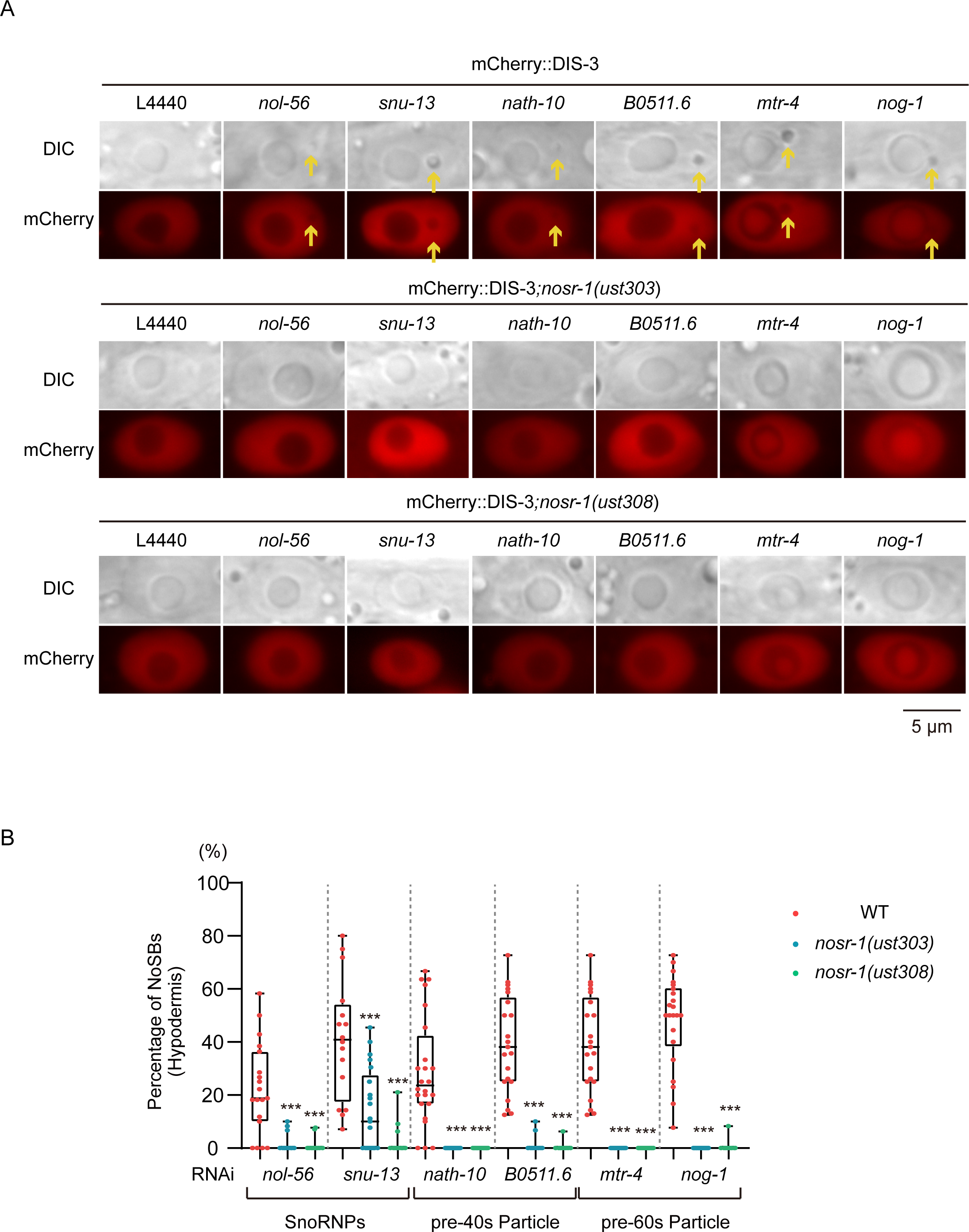
NOSR-1 is required for NoSB formation. (A) DIC and fluorescence microscopy images of *C. elegans* nuclei after knocking down indicated genes. (B) Quantification of NoSB in hypodermic cells from (A). Every dot indicates the percentage of hypodermis cells containing NoSB in each worm. n>19 animals.

**Figure S10.**
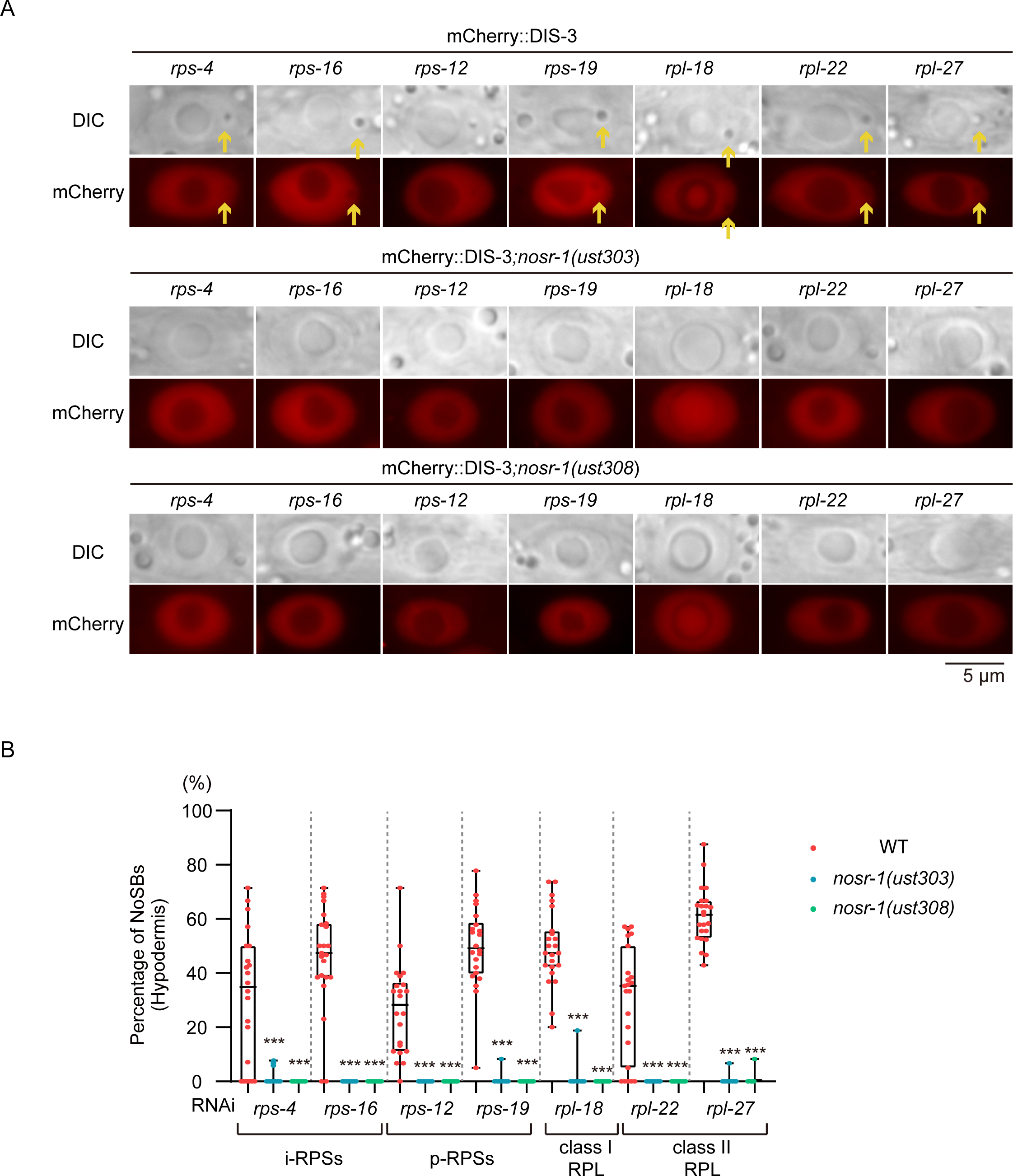
NOSR-1 is required for NoSB formation. (A) DIC and fluorescence microscopy images of *C. elegans* nuclei after knocking down indicated genes. (B) Quantification of NoSB in hypodermic cells from (A). n>19 animals.

**Figure S11.**
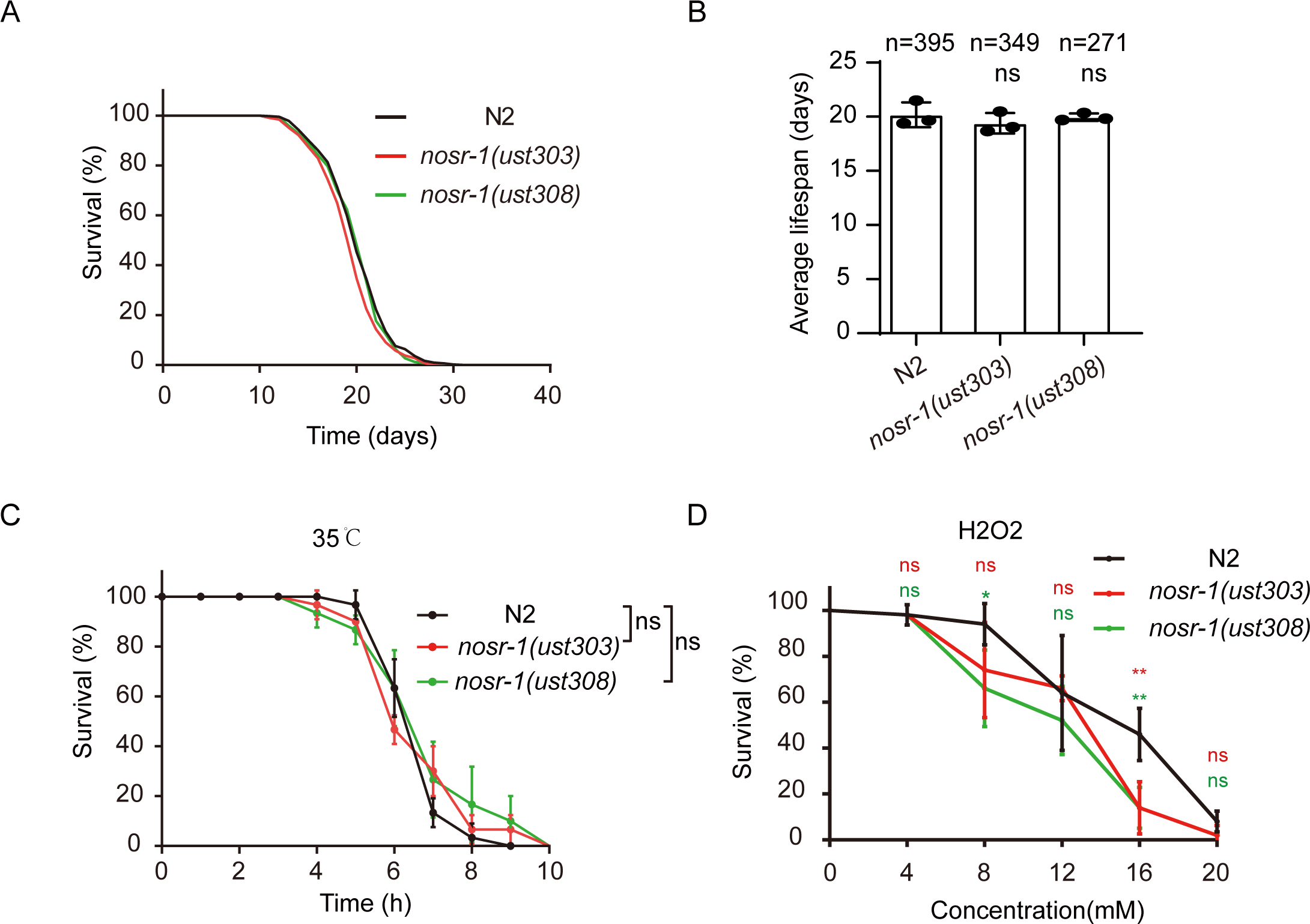
The effects of *nosr-1* mutation on lifespan and anti-stress response. (A) Survival curves of the indicated animals at 20°C. The same lifespan data of control N2 worms were used in Figures 5E-F. (B) Histogram displaying the average lifespan of the indicated animals. Mean ± SD of three independent experiments. Asterisks indicate significant differences using log-rank tests. ns, not significant, p>0.05. (C) Heat stress survival curves of the indicated animals. Data in each panel are presented as the mean ± SD of three independent experiments. (D) Oxidative survival curves of the indicated animals.

**Figure S12.**
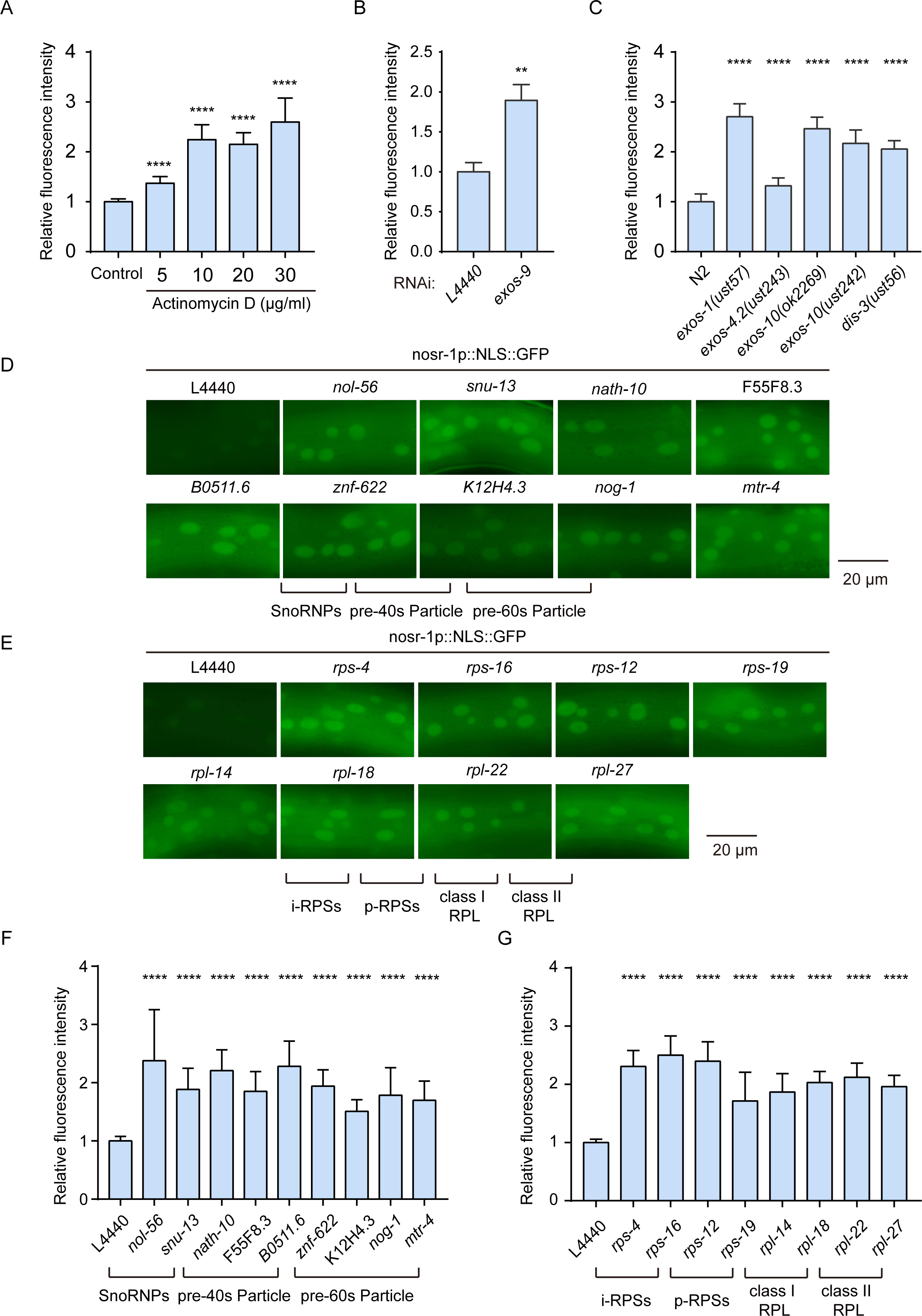
Deficient rRNA biogenesis activates *nosr-1* expression. (A, B, C) Quantification of fluorescence intensity in **Figs. 6B-D**, respectively. (D, E) Fluorescence microscopy images of *nosr-1p::NLS::GFP* after knocking down the indicated genes by RNAi. (F, G) Quantification of fluorescence intensity in **panels D-E**, respectively. Data are presented as the means ± SD of at least 20 worms for each worm strain. ****P < 0.0001.

**Figure S13.**
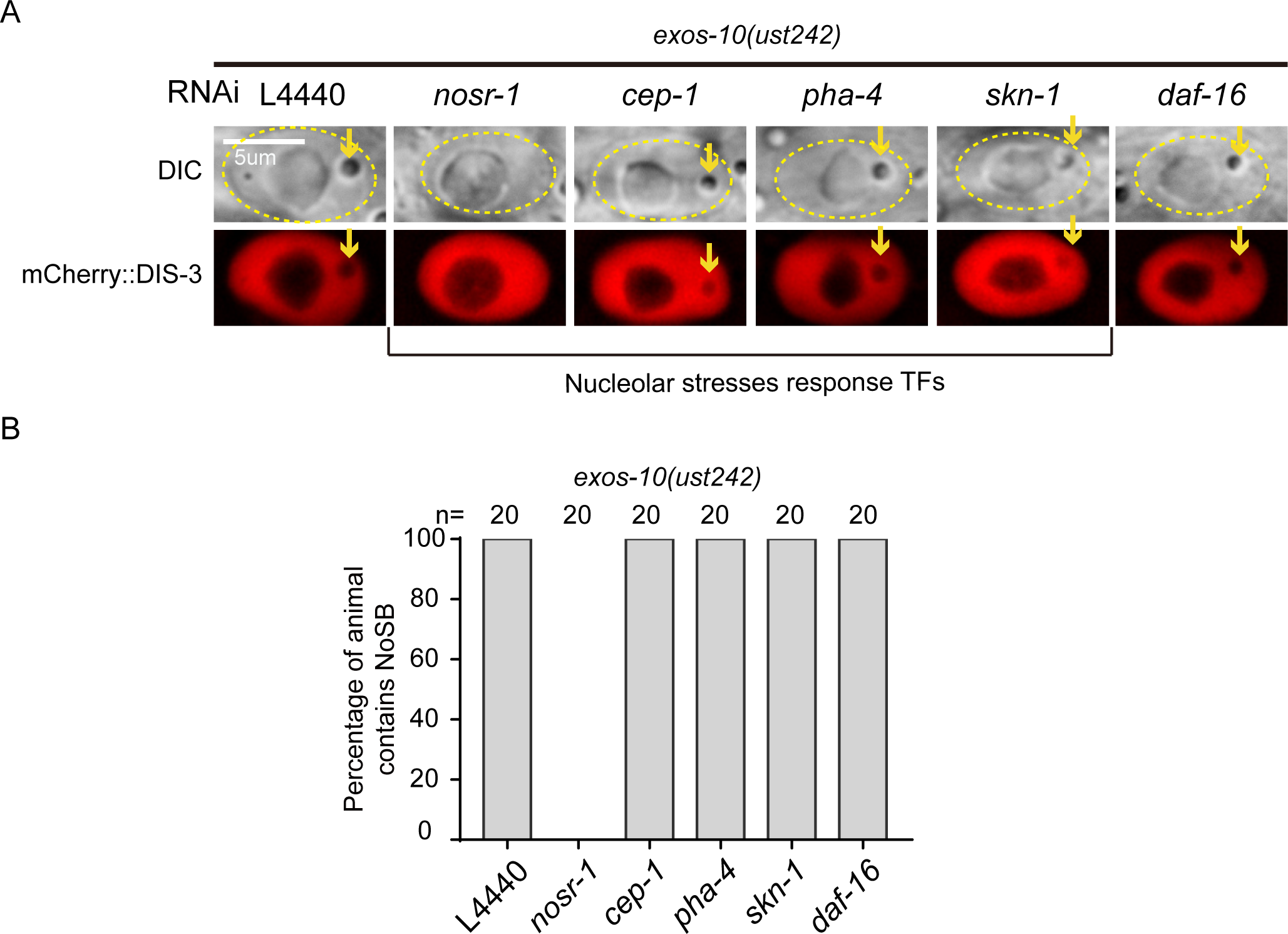
*nosr-1* may represent a new nucleolar stress responsive pathway. (A) DIC and fluorescence microscopy images of hypodermal nuclei in *exos-10(ust242);mCherry::dis-3* worms after knockdown of the indicated genes by RNAi. (B) Quantification of NoSB in hypodermic cells in the indicated animals. The percentage of animals containing at least three NoSB structure were shown. n=20 animals.

**Figure S14.**
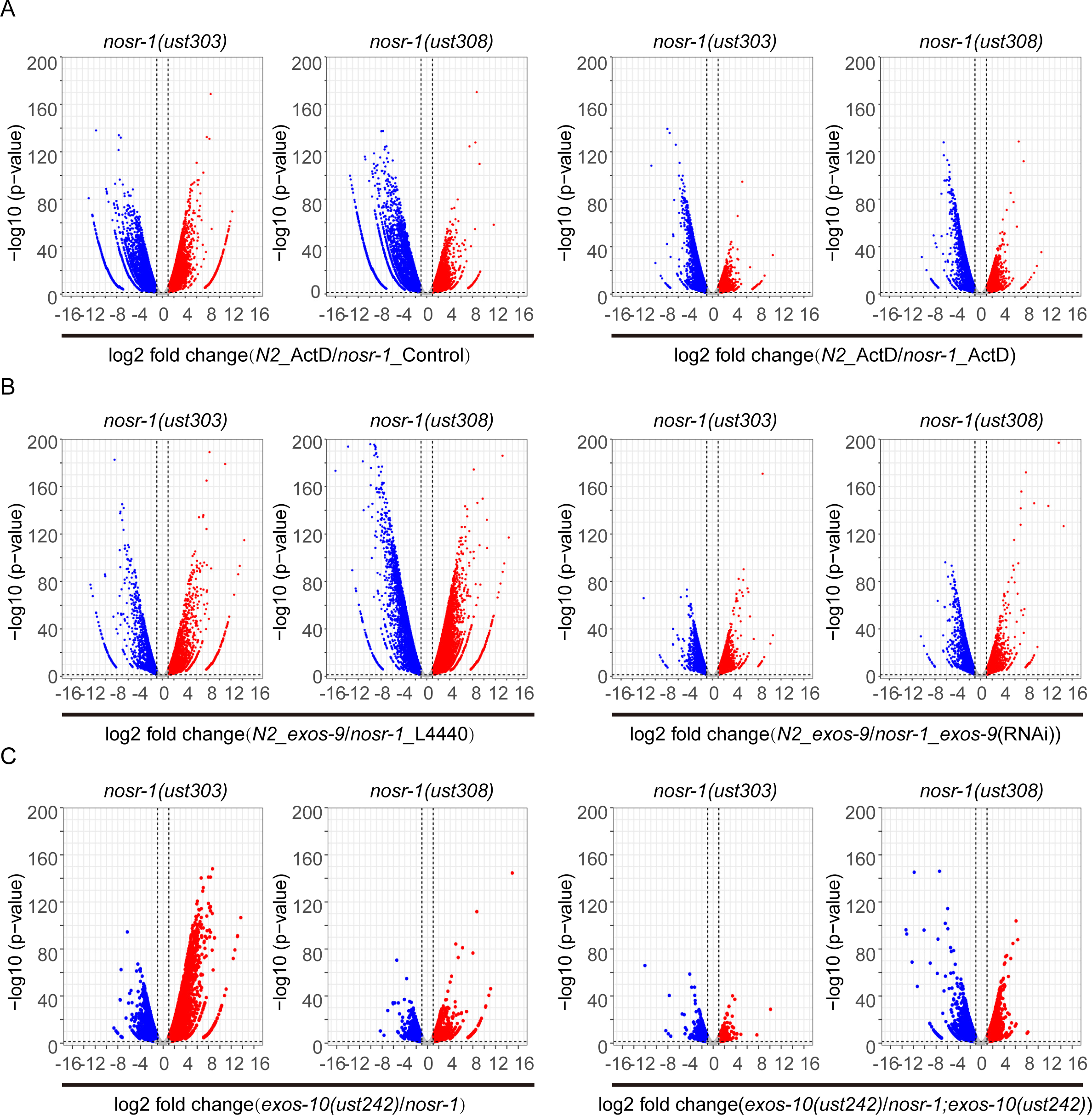
mRNA-seq identified NOSR-1 dependent, nucleolar stress-induced differentially expressed genes. (A-C) Volcano plot comparing gene expression between the indicated animals. The upregulated genes were defined as those with a fold change ≥ 2. The downregulated genes were defined as those with a fold change ≤ 0.5.

**Figure S15.**
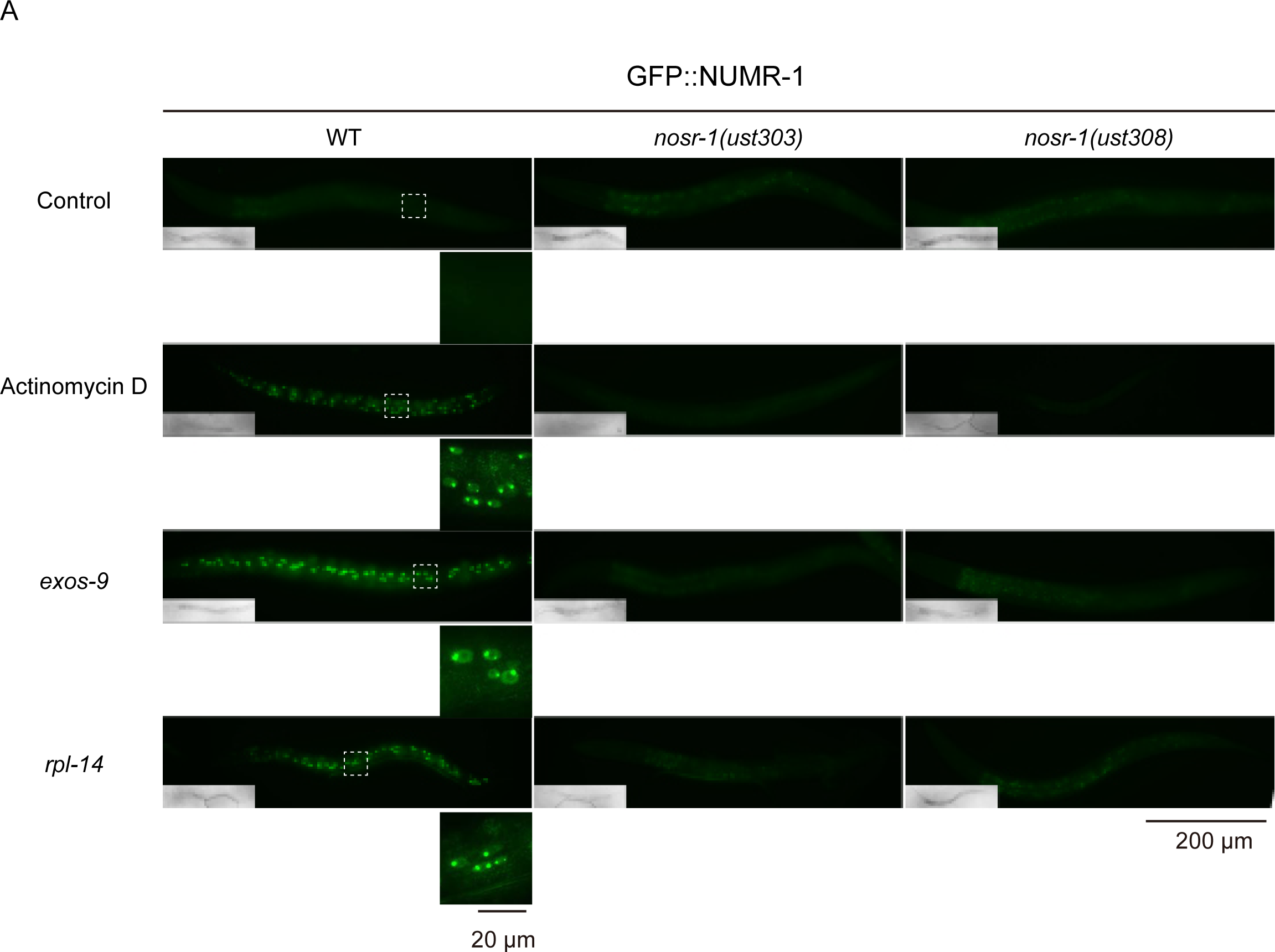
Nucleolar stress activates *numr-1* expression via *nosr-1*. (A) Fluorescence microscopy of L3/L4 stage animals after actinomycin D treatment and RNAi knockdown of *exos-9* and *rpl-14*, respectively, in the indicated animals.

**Figure S16.**
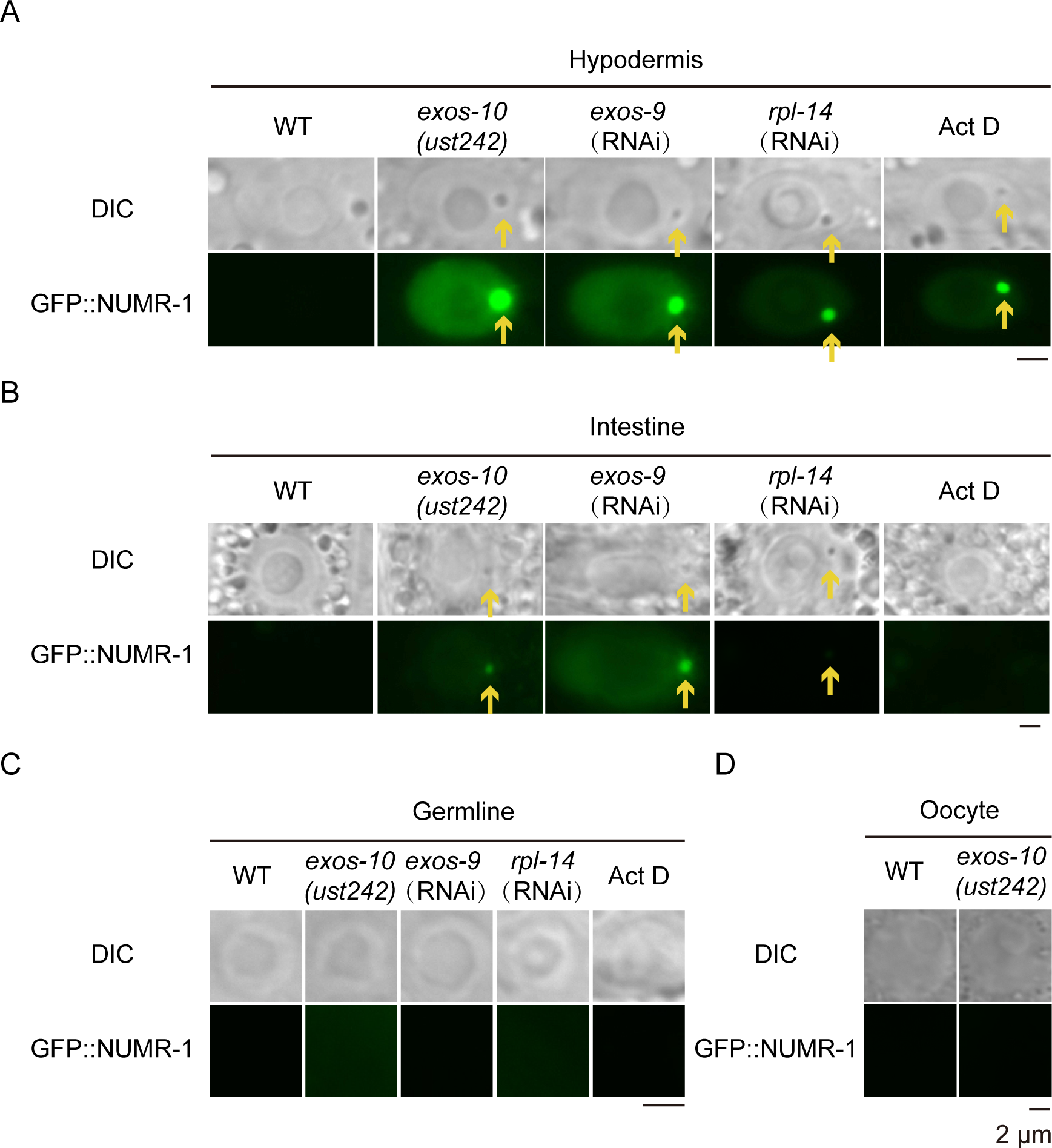
Tissue specific formation of NoSB. (A-B) DIC and fluorescence microscopy images of nuclei in indicated tissues after knockdown of the indicated genes or Actinomycin D treatment in the indicated animals. (C-D) DIC images of nuclei in indicated tissues after knockdown of the indicated genes or Actinomycin D treatment in the indicated animals.

**Figure S17.**
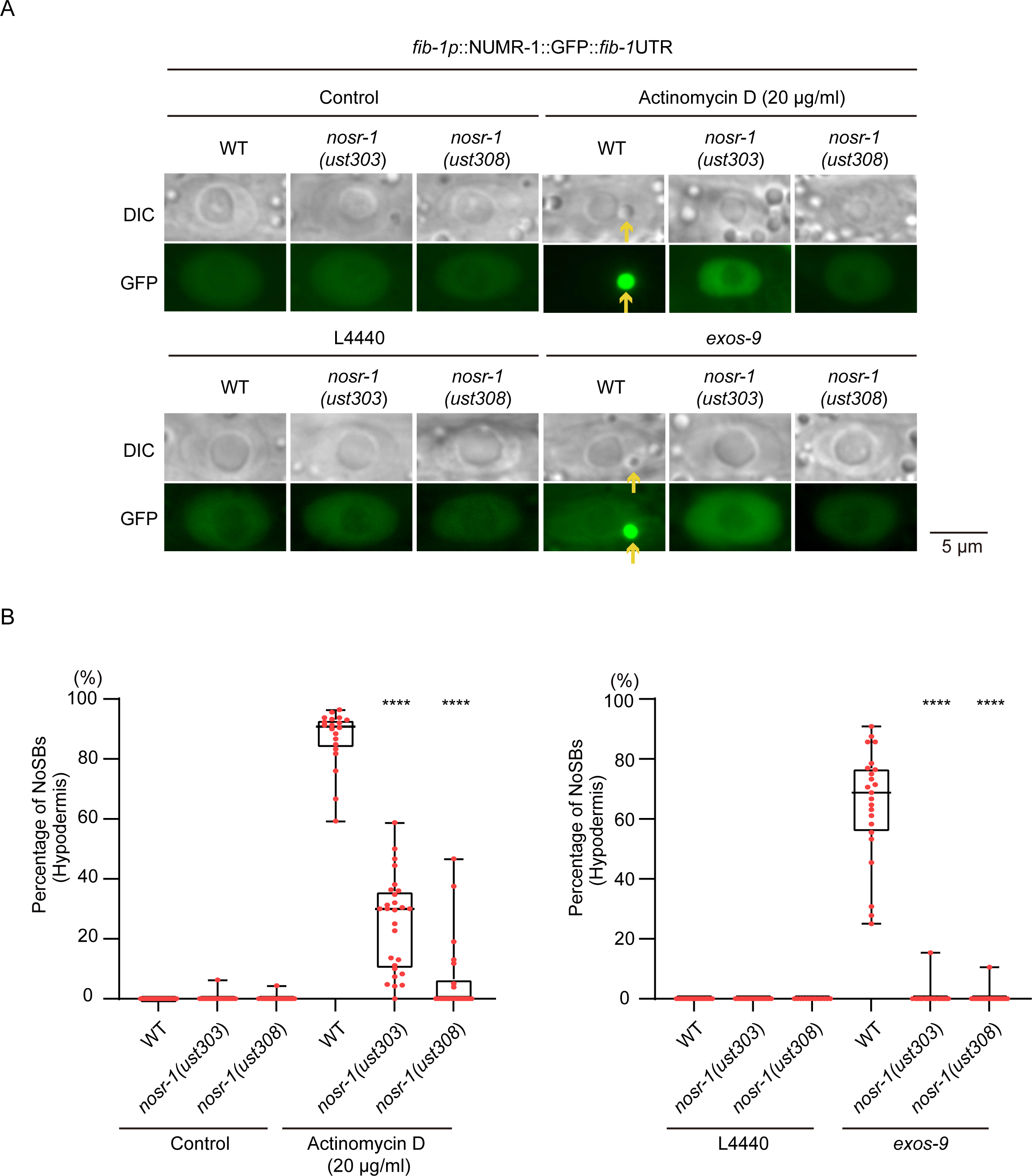
Nucleolar stress induced a NOSR-1-dependent condensation of NUMR-1::GFP. (A) DIC and fluorescence microscopy images of nuclei in indicated tissues after knockdown *exos-9* or Actinomycin D treatment in the indicated animals. (B) Quantification of the percentage of NoSB in **panel A**. Every dot indicates the percentage of hypodermis cells containing NoSB in each worm. n>19 animals.

**Figure S18.**
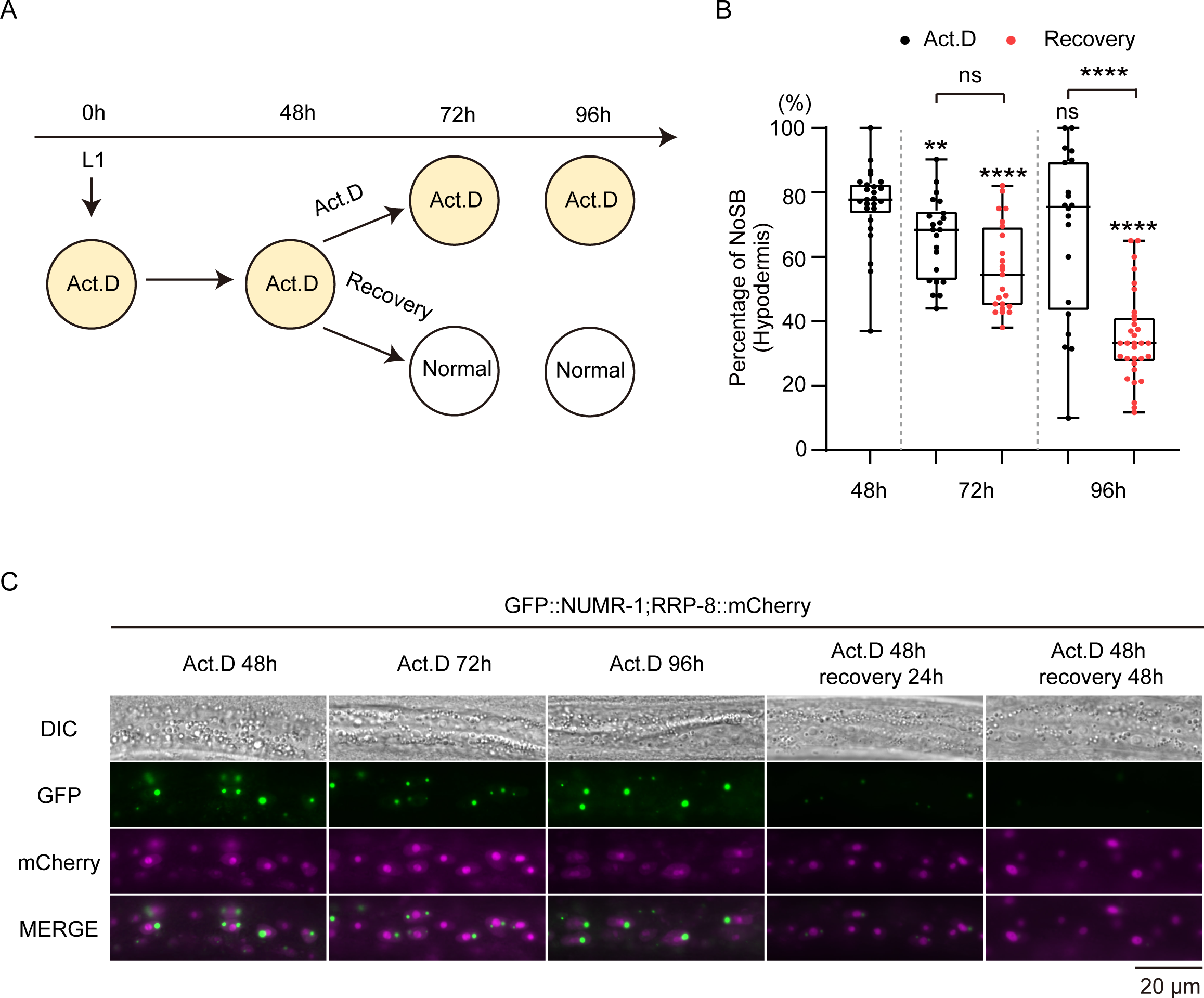
NoSB disassembled upon the removal of nucleolar stress. (A) Schematic diagram of nucleolar stress recovery assay. (B) Quantification of the percentage of NoSB upon the removal of actinomycin D. Every dot indicates the percentage of hypodermis cells containing NoSB in each worm. n>19 animals. (C) Fluorescence microscopy images showing epidermal cells of the indicated animals expressing GFP::NUMR-1 and RRP-8::mCherry upon the removal of actinomycin D.

**Figure S19.**
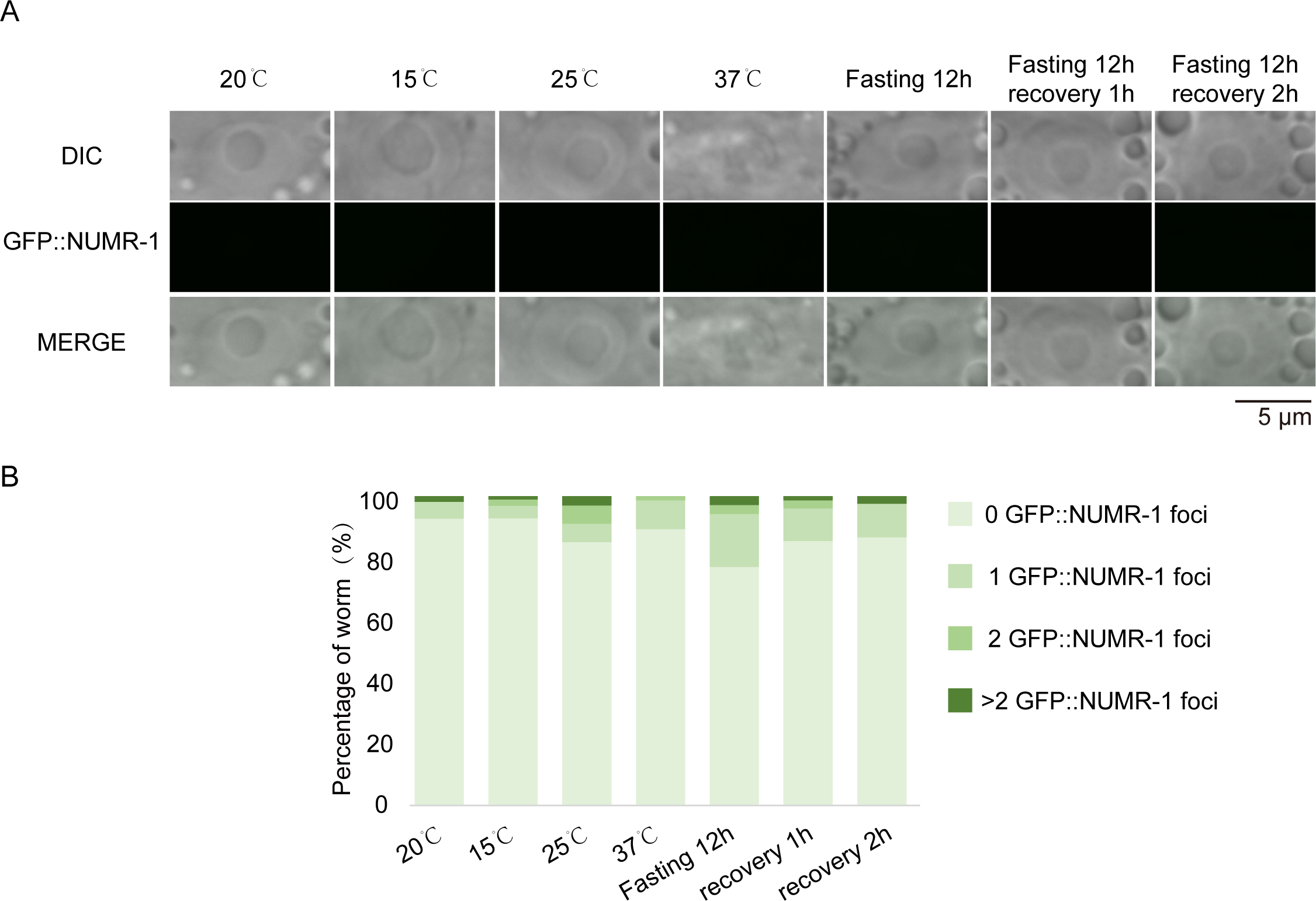
Temperate shift did not induce NoSB formation. (A) DIC and fluorescence microscopy images of nuclei at the indicated temperatures conditions. (B) Quantification of the percentage of worms with indicated number of GFP::NUMR-1 foci in each cell.

**Figure S20.**
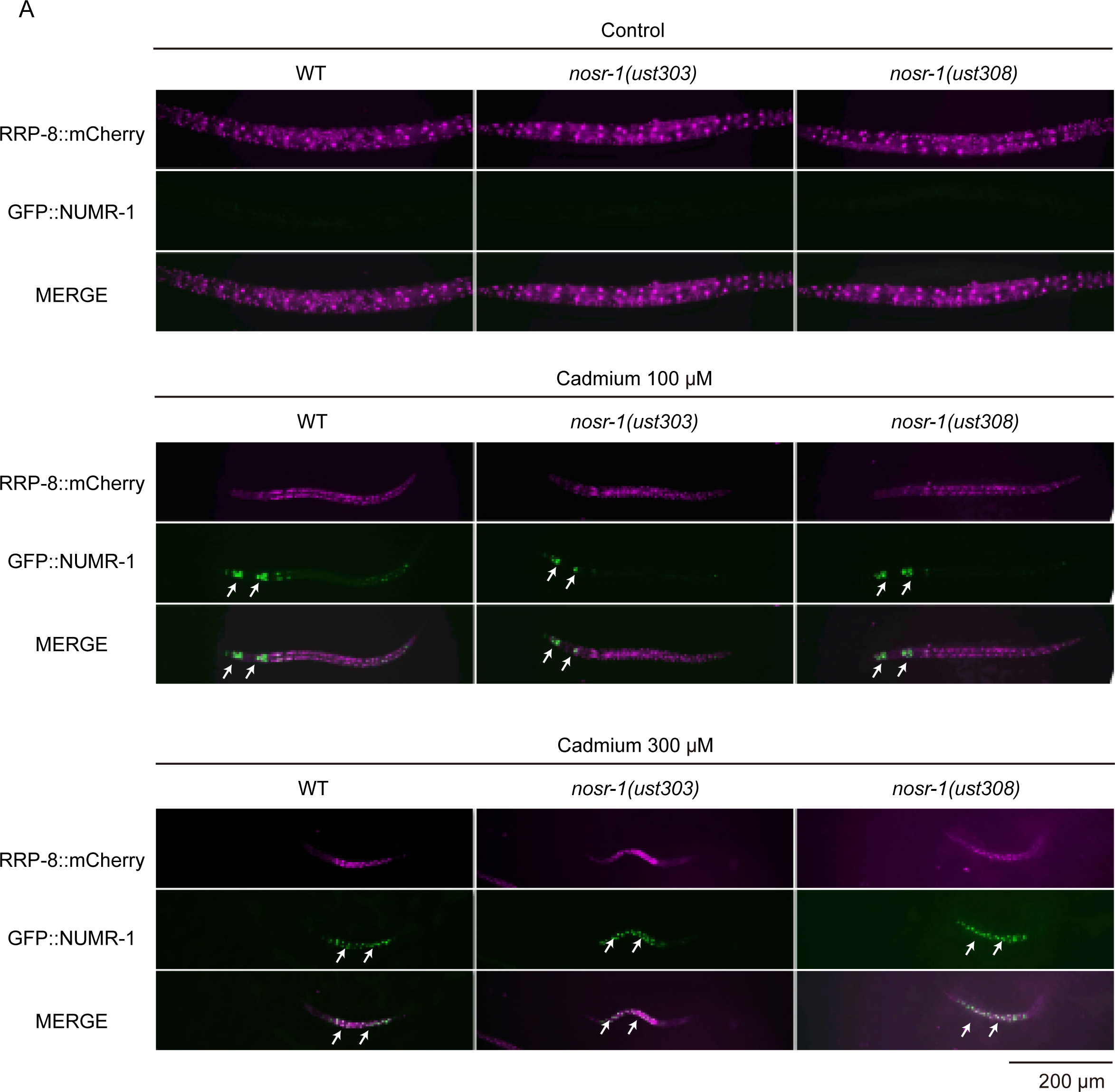
Cadmium stress induced GFP::NUMR-1 expression. (A) Fluorescence microscopy images of L3/L4 stage animals after cadmium treatment in the indicated animals.

**Table S1.**
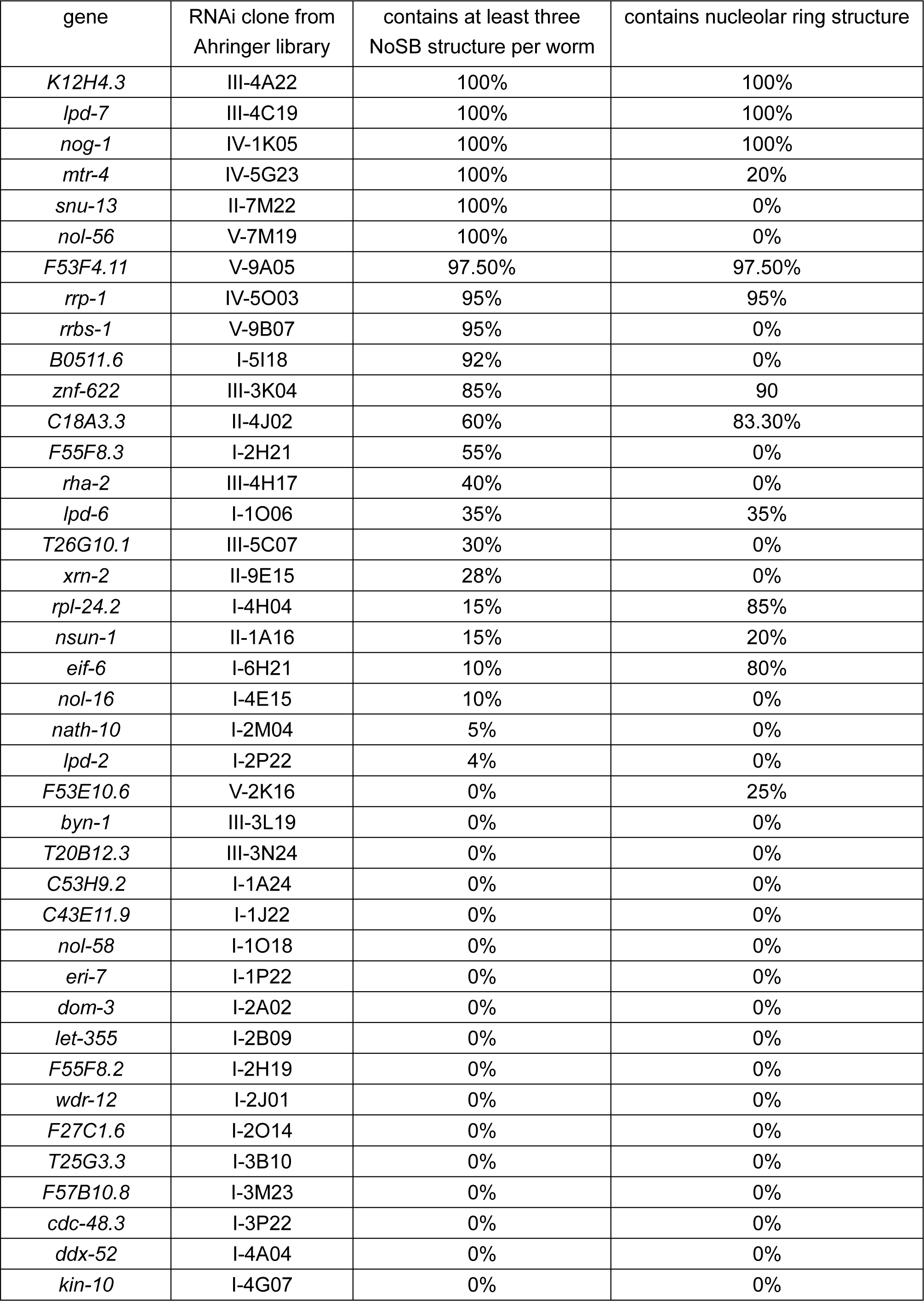
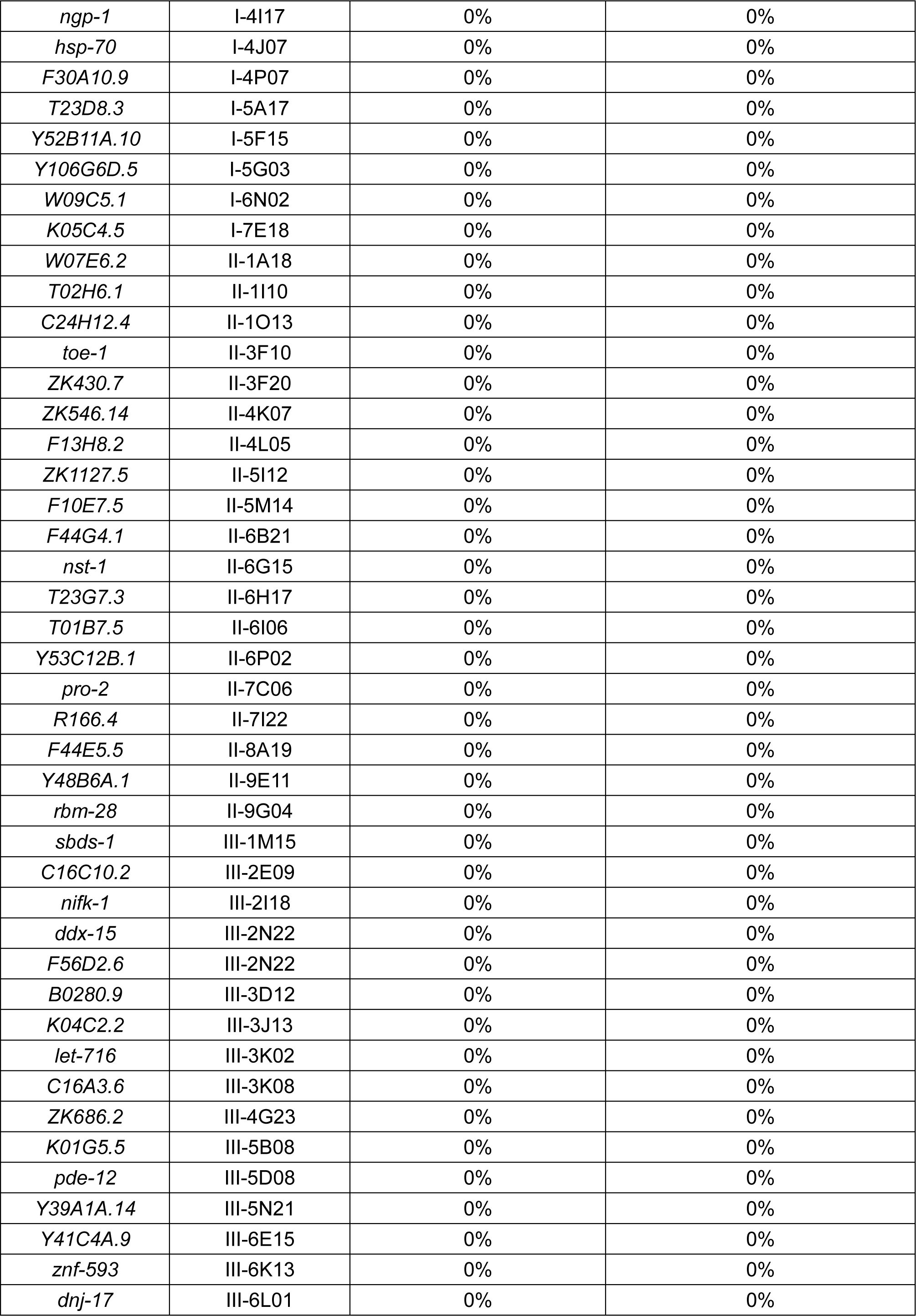
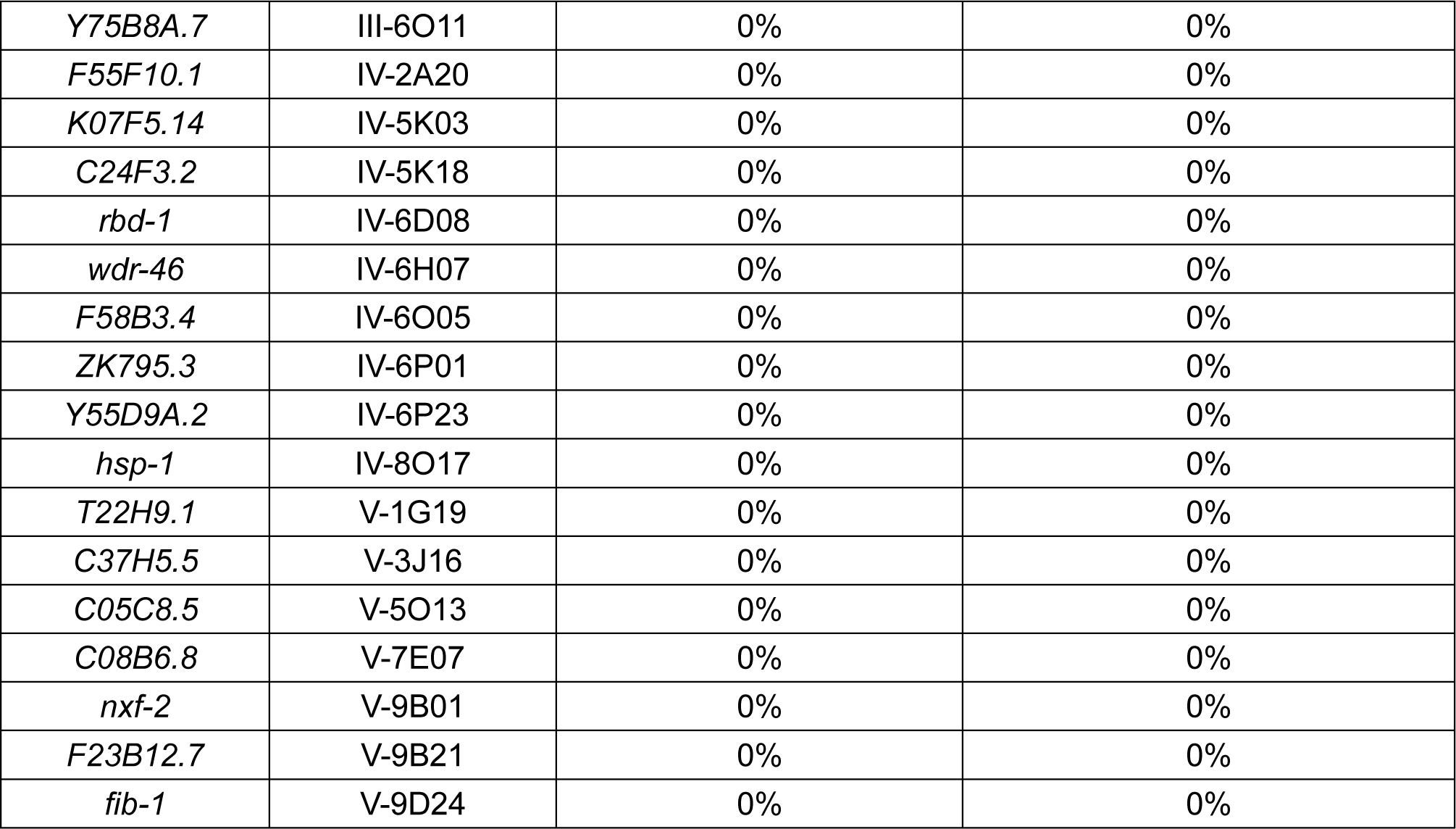
List of genes used in the candidate-based RNAi screening.

**Table S2:**
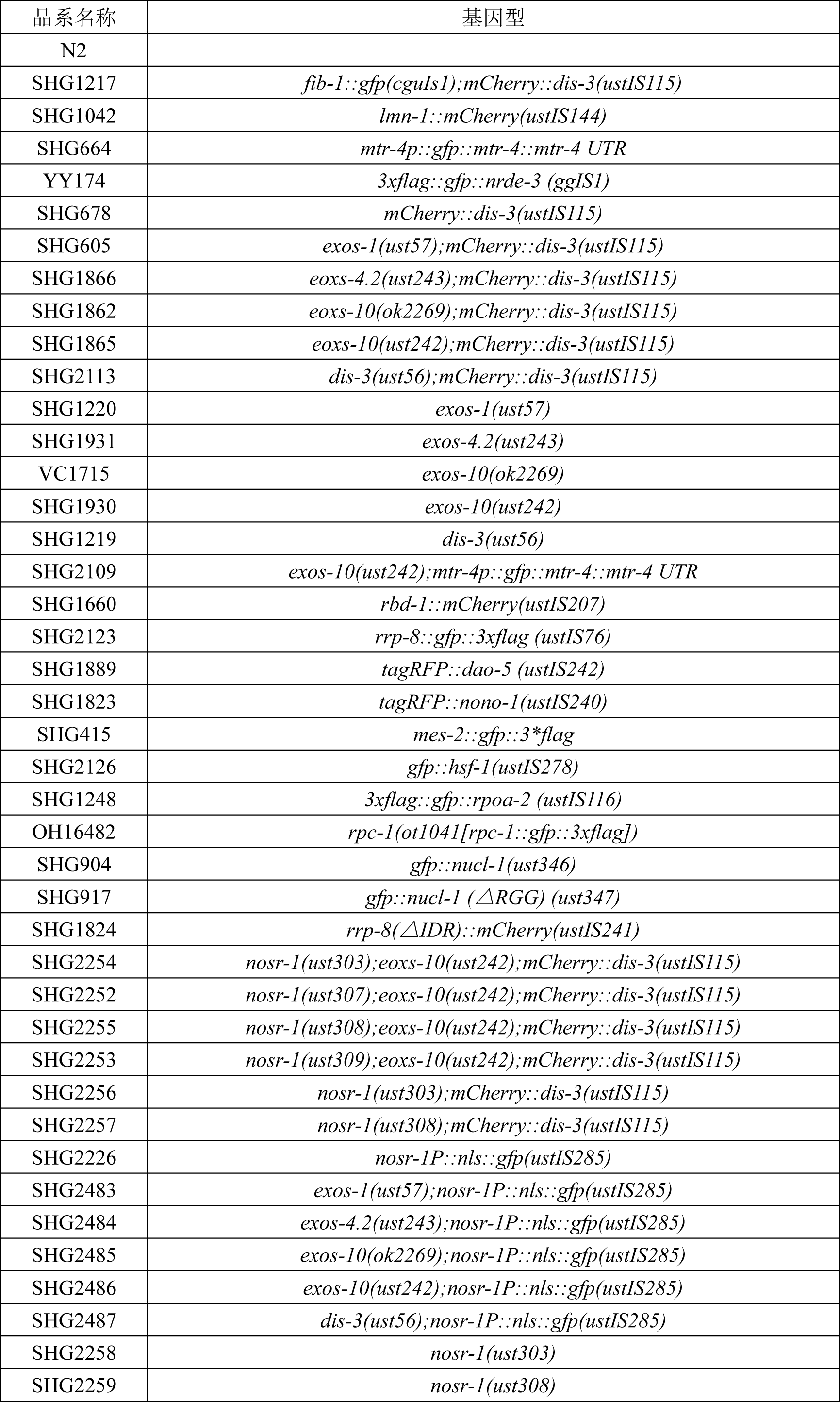
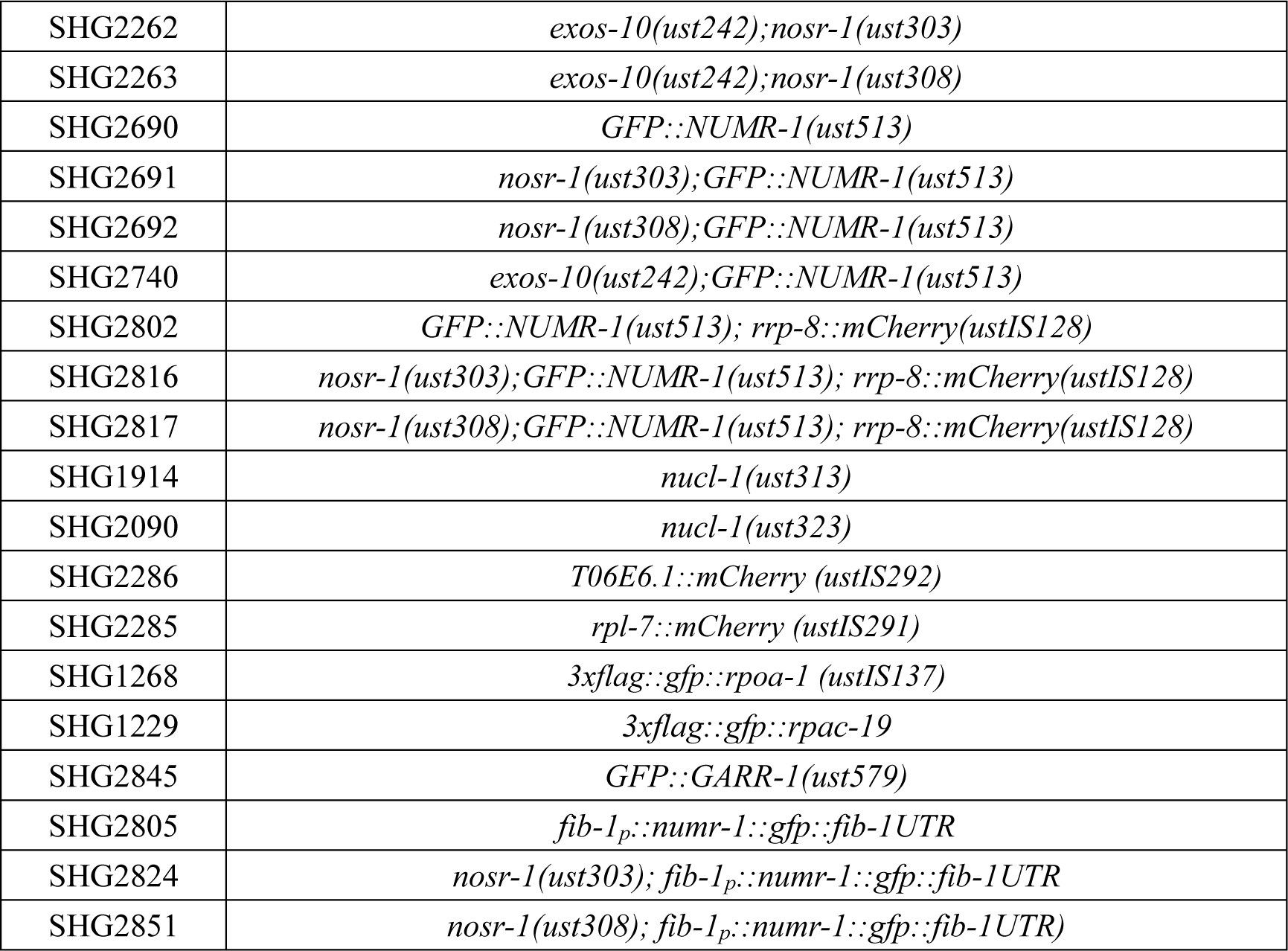
List of strains used in the work.

**Table S3:**
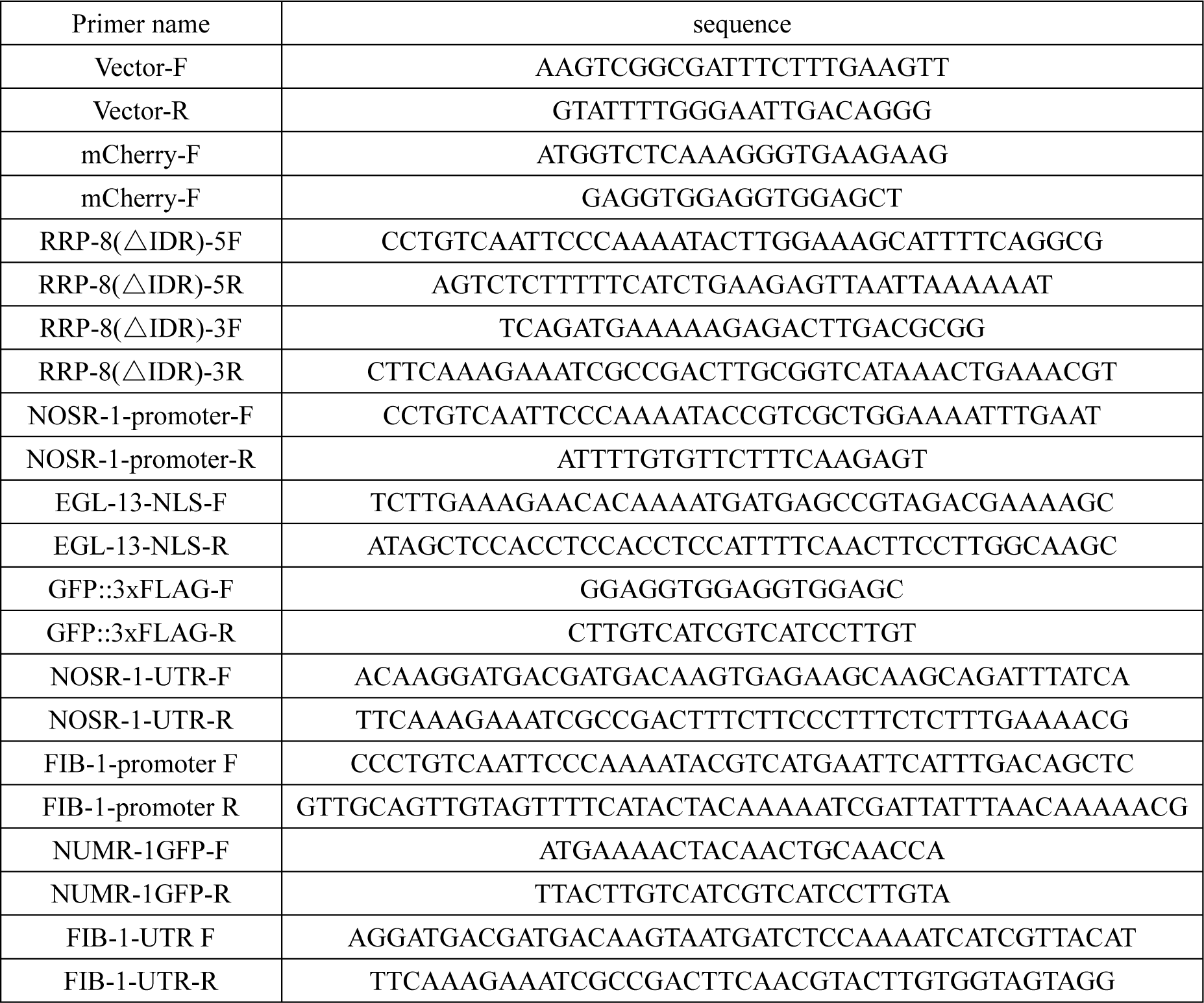
Sequence of repair plasmid primers used in ectopic transgenic strain construction.

**Table S4:**
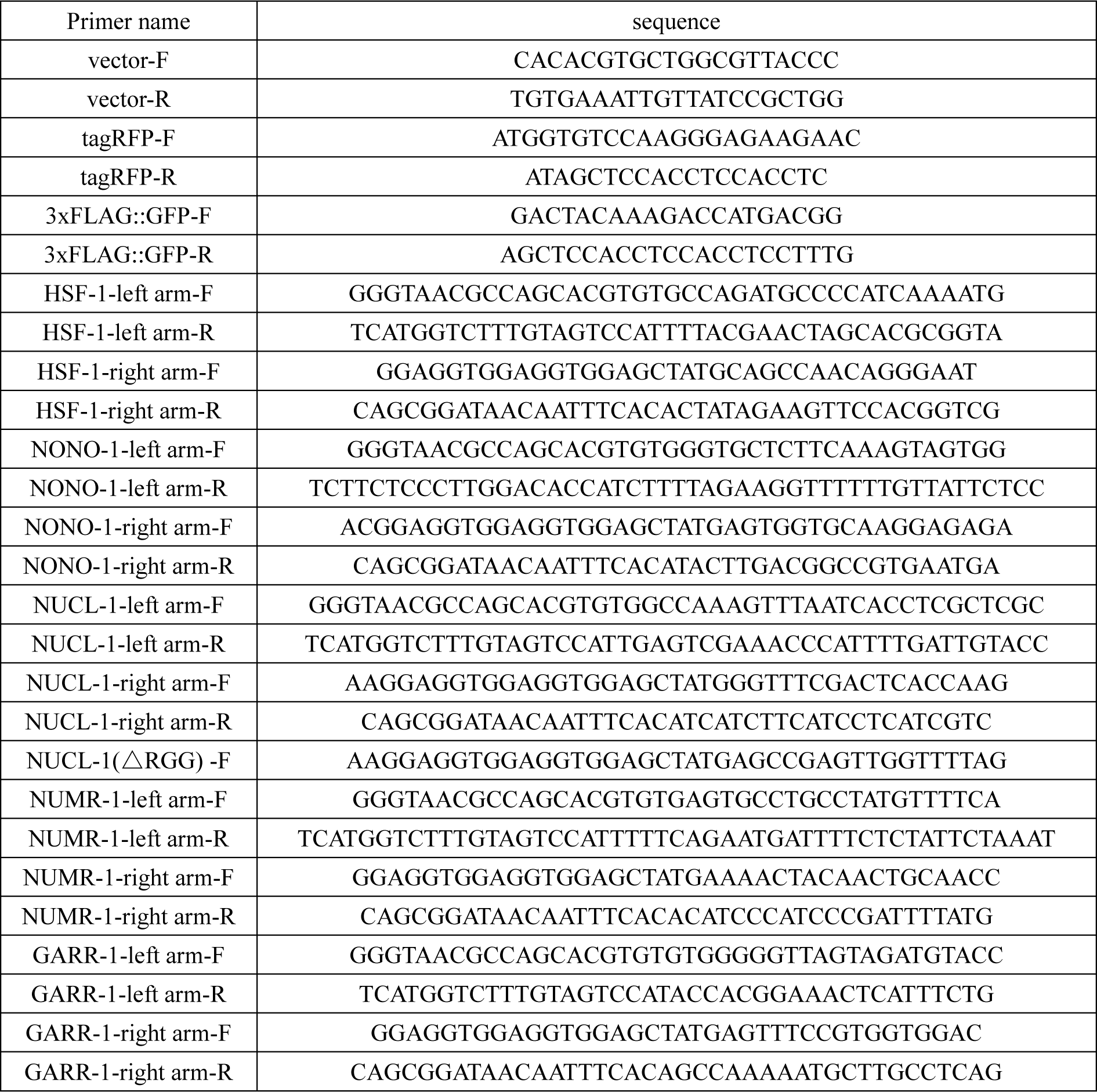
Sequence of repair plasmid primers used in in situ transgenic strain construction.

**Table S5:**
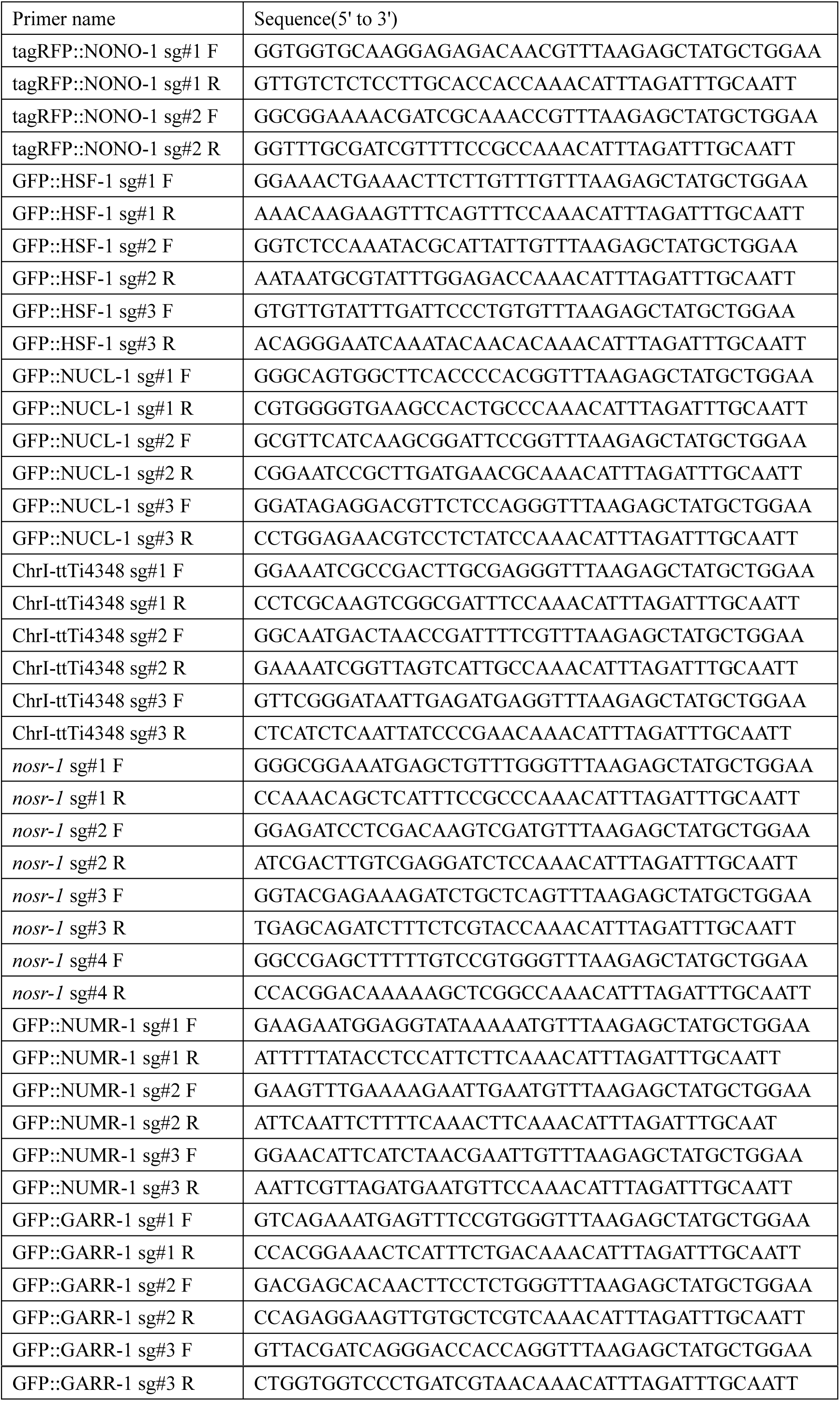
Sequences of sgRNAs for CRISPR Cas9-mediated gene editing.

**Table S6:**
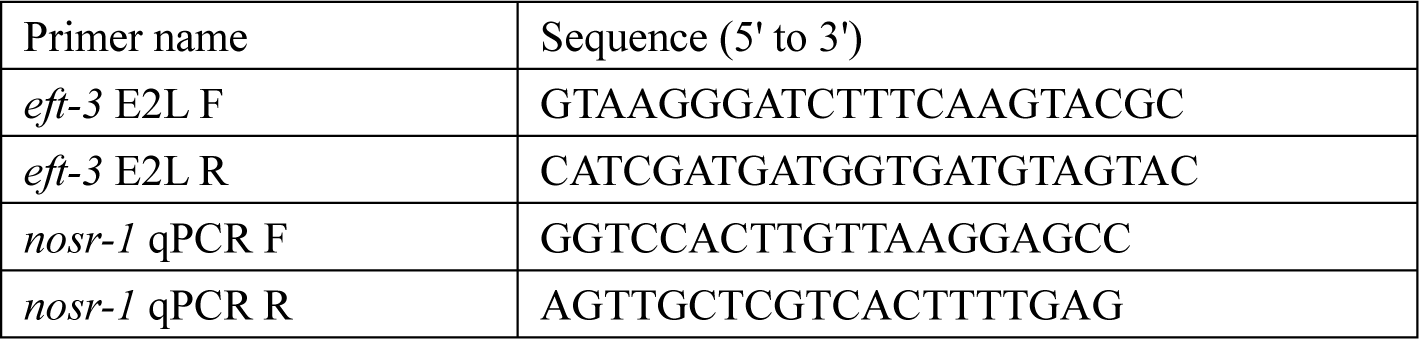
Sequences of quantitative real-time PCR primers.

